# *intI*1 primer selection for class 1 integron integrase gene and transcript quantification – validation and application for monitoring *intl*1 gene abundance within septic tanks in Thailand

**DOI:** 10.1101/2023.06.19.545554

**Authors:** Valentine Okonkwo, Fabien Cholet, Umer Z. Ijaz, Thammarat Koottatep, Tatchai Pussayanavin, Chongrak Polpraset, William T. Sloan, Stephanie Connelly, Cindy J. Smith

**Affiliations:** Infrastructure and Environment, James Watt School of Engineering, University of Glasgow, UK; School of Environment, Resources and Development, Asian Institute of Technology, Thailand; Faculty of Science, Ramkhamhaeng University, Thailand; Thammasat School of Engineering, Thammasat University, Thailand

**Keywords:** Antimicrobial resistance (AMR), Antibiotic resistance genes (ARGs), Class 1 integron-integrase (*intl*1) gene, septic tank, (RT)-QPCR, MiSeq, mRNA transcripts

## Abstract

Antimicrobial resistance (AMR) poses serious global public health threat and wastewater treatment (WWT), including septic tanks, are a significant source of AMR genes to the environment. Environmental monitoring of broad-range AMR genes remains a challenge. The class-1 integron-integrase (*intI*1) gene has been proposed as a proxy for overall AMR abundance, but there is no consensus on the qPCR primer set to use. A systematic review of the literature found 65 primer sets. The coverage and specificity of each, including newly designed MGB-*Taq*Man primer-probe, was evaluated against a database of clinical and environmental *intl*1, intl1-like and non-intl1 sequences. Three primer sets were selected, laboratory validated for DNA and mRNA quantification and used to quantity *intl*1 gene abundance from household and healthcare conventional septic tanks (CST) and novel household Solar Septic Tanks (SST) in Thailand. Specificity of the *intl1* septic tank amplicons showed that no primer set could distinguish between *intl1* and *intl-1* like sequences. Each primer set showed the same trends across septic tanks, with highest gene abundance in influent>sludge>effluent. There was no statistical difference between the same sample quantified by the three primer sets. However, when comparing gene abundances from the same primer set across septic tanks, statistical differences between different sample types were observed for one primer set but not the others. This may lead to different interpretation of risk associated with each reactor in spreading *intl*1 to the environment. Comparing reactor types*, intI*1 abundance in the effluent was lowest in the SST-household<CST-household<CST-healthcare. Depending on primer set used, 31 to 42% of *intI*1 was removed from effluent of the CST-household tank with accessible influent. Our study provided insight into the importance of *intl1* primer choice. We propose the use of the validated set (F3-R3) for optimal *intI*1 quantification and towards the goal of achieving standardisation across environmental studies.

## Introduction

Antimicrobial resistance (AMR), the ability of microbes to grow and thrive in the presence of compounds capable of limiting their cellular growth or kill cells, is a serious growing public health concern globally; and has recently been classified as “*one of the top ten global threats facing humanity*” by the World Health Organisation (WHO, 2021).

The occurrence of AMR via mutation and subsequent vertical gene transfer or acquisition of AMR genes via horizontal gene transfer (HGT), is an inevitable natural phenomenon in the evolution of microbes (Holmes et al., 2016; Hayward et al., 2019). Nonetheless, recent global challenges including extensive consumption and misuse of antimicrobials, particularly antibiotics, in clinical settings, agri-and aqua-culture and their subsequent release to the environment, have given rise to the emergence and rapid dissemination of AMR genes amongst bacteria, including microbes of clinical importance, and to the environment (Holmes et al., 2016). Consequently, high global mortality, as a result of patient treatment failure, has been associated with AMR-related infections (700, 000 deaths in 2014) (O’ Neill, 2014). Moreover, the global death toll from AMR-related infections has been projected to increase to 10 million deaths per year by 2050 surpassing death from cancer, assuming no change to the current trends/policies, coupled with an economic burden of 100 trillion US Dollars (O’ Neill, 2014).

Wastewater treatment (WWT), including decentralised treatment systems such as septic tanks, receive significant amounts of antibiotics from human and animal waste (30 to 90% of antibiotics are excreted in urine and faeces (Sarmah et al., 2006)) and are now recognised as important reservoirs for AMR creating hotspots for transfer and subsequent release to the environment (Rizzo et al., 2013; Connelly et al., 2019; Hayward et al., 2019). The selective pressure introduced by the often multiple, low-levels, sub-inhibitory concentrations of antimicrobials found in wastewater (WW), promotes AMR gene acquisition amongst microbes via HGT and selection for AMR bacteria. WWT and septic tanks are unable to effectively remove these (Gillings et al., 2015; Gillings, 2017; Hayward et al., 2019), resulting in increased AMR genes and bacteria discharged directly to the environment, contributing significantly to the global burden (Amos et al., 2018). The global AMR burden from wastewater is further exacerbated in the Global South due to high prevalence of extensive antibiotic usage-propelled by poor regulations on usage, ineffective or lacking WWT, coupled to increasing populations and rapidly expanding megacities.

The necessity to tackle AMR discharge from WWT to the environment requires a comprehensive understanding of the role of WWT in the dissemination of AMR to the environment. This understanding will create unique opportunities to implement key strategies to mitigate AMR spread, and in turn, allows for the safeguarding of global public health. This knowledge can be informed by sensitive, accurate detection, quantification, and tracking of AMR genes from source (e.g. WWT) to the environment. However, multiple AMR genes exist within WWT. Monitoring numerous AMR genes simultaneously is a major challenge (Gillings et al., 2015), particularly if a rapid assessment is needed. Similarly, monitoring one or a subset of AMR genes is not ideal, as selected AMR gene(s) may be absent (Gillings et al., 2015). Previously, the clinical class 1 integron (CL1-integron) integrase (*intI*1) was proposed as a proxy for inferring potential AMR, which circumvents multiple monitoring limitations, by acting as proxy for potential AMR pollution (Gillings et al., 2015). *Intl*1 was proposed as a proxy as: it is linked to genes that confer resistance to antibiotics, disinfectants and heavy metals; it is found in diverse taxonomic groups of pathogenic and non-pathogenic bacteria and can move across taxa via HGT due to its physical linkage to mobile genetic elements (MGEs) such as plasmid and transposons; its abundance can rapidly change in response to external pressures such as the presence of antibiotics; selection pressures imposed by recent human activities have resulted in the emergence of the highly conserved clinical *intI*1 variant (Gillings et al., 2015), the elevated presence of which in the environment indicates pollution and potential hotspot for AMR transfer (Gillings et al., 2015; Pruden et al., 2021).

Currently, molecular approaches, specifically real-time quantitative PCR (Q-PCR), have emerged as the methods of choice for AMR and CL1-integron detection and quantification in the environment. By far the most prevalent approach for detecting or quantifying the CL1-integron is the amplification of the *intI*1 gene at the 5’ conserved segment (CS) across diverse ecological niches including engineered systems e.g. WWTs (Chen and Zhang, 2013; Berglund et al., 2015; Li et al., 2016) and natural ecosystems such as sediments (Lapara et al., 2011; Dong et al., 2019). Whilst targeting the *intI*1 gene provides no information about the structure beyond the 5’ CS, quantification of the *intI*1 gene as an initial screening to infer potential AMR contamination within complex environments is invaluable and a useful initial screening approach. However, within the literature numerous primers targeting the *intI*1 gene are available (**see Table S.1**) and different sets are used across different studies. The current lack of standardisation prevents cross study comparisons and limits current understanding of AMR in the environment. As such, selecting an optimal *intl*1 primers with both high coverage and specificity suitable for environmental monitoring is a challenge. Moreover, several primers have been designed based on the highly conserved clinical *intl*1 gene sequences (≥98% protein similarity to each other), and the extent to which these primers target the less conserved *intI*1 gene variants (<98% protein similarity) found also in environmental samples (Gillings et al., 2008a, 2008b; Gillings et al., 2015) and on the chromosome non-pathogenic *Betaproteobacteria* which carries gene cassettes not associated with AMR genes (Gillings, et al., 2008a), has yet to be determined. As such a comprehensive and comparative evaluation of published *intI*1 primers to determine their coverage and specificity against clinical and environmental *intl*1 sequences to identify a consensus optimal *intI*1 primers for monitoring AMR within environmental samples is urgently needed (Zhang et al., 2018).

With this need identified, we undertook to review, evaluate, and then apply *intI*1 primers to quantify the gene across a suite of wastewater samples from septic tanks in Thailand. Specifically, we compare the recent solar septic tank (SST) technology currently implemented in some areas of Thailand (Polprasert et al., 2018; Connelly et al., 2019) to that of conventional septic tanks (CST) treating household and healthcare wastewater. The SST technology differs from CST primarily by the incorporation of a central disinfection chamber containing a heated copper coil connected to a passive solar heat collection system installed on the roof of the toilet block served by the SST (Polprasert et al., 2018; Connelly et al., 2019). The heat from the central chamber (50 - 60°C by design) promotes partial pasteurisation as the effluent passes through the chamber prior to discharge. Effluent water quality is improved by reducing microbial biomass including potential pathogens, and by extension reduction of the microbial load should reduce the AMR burden to receiving water bodies. Moreover, the in-tank temperature is raised by the passive transfer of heat from the chamber to the rest of the tank; thus, promoting enhanced microbial degradation of both retained solids (sludge) and soluble compounds (Connelly et al., 2019). As such, we hypothesis that *intl*1 gene abundance would be lower in the SST than the CST sludge and effluent owing to the enhanced treatment caused by the increased temperature.

To address this methodological knowledge gap and our hypothesis a systematic review of the literature was undertaken to obtain published *intI*1 primers followed by a comprehensive *in-silico* analysis of primer coverage and specificity against a curated database of clinical and environmental *intl*1 sequences to select the best performing primers. A subset of the best performing primer sets was used to quantify *intI*1 gene abundance from 30 septic tank wastewater samples comparing conventional (healthcare and household wastewater) and solar septic tank (household wastewater), with *intI*1 specificity validated by Illumina MiSeq. We further confirmed the suitability of the primers to quantify *intI*1 gene transcripts. Thus, we propose validated *intl*1 primer set for quantification of genes and transcripts from environmental samples towards the goal of achieving standardisation across *intI*1 studies.

## 2. Materials and Methods

### 2.1 *intI*1 primer evaluation

#### 2.1.1 Systematic review of literature and alignment of primers to *intI*1 reference sequence

A systematic review of >3000 peer-reviewed publications was conducted to retrieve *intI*1 primers and probes sequences across a range of settings including clinical and environmental e.g., agricultural, and human-impacted settings including WWTPs. For this, the “Web of Knowledge” database (https://www.webofscience.com/; last assessed 04/10/2022) was searched using the term “Class 1 integron”. Only published articles in English language were considered. 3266 published articles were subsequently recovered. The *intI*1 primer sequences from the respective literature were either retrieved in the main text or from the accompanying supplementary material.

Obtained *intI*1 primer and probe sequences were aligned to a reference *Pseudomonas aeruginosa* plasmid pVS1 nucleotide sequence (M73819.1) using the ClustalX2 algorithm (Version 2.1.0.0), with default settings (Larkin et al., 2007) and visualised with BioEdit (version 7.0.5.3) (Hall, 1999). The alignment position of each primer and probe sequence were renamed according to position along the *Pseudomonas aeruginosa* reference *intI*1 gene sequence (Figure S.1, Table S.1).

#### 2.1.2 Databases construction and curation

The integron-integrase database by Zhang et al., (2018) consisting of 922 and 2462 *intI*1 gene and integron-integrase (*intI*) of other class protein sequences respectively (herein referred to as non-*intI*1 database) (Figure 1) was employed for the analysis of primers. Whilst the *intI* of other class database was mostly populated with protein sequences from other integron-integrase classes, it also contained a number of XerCs integrases (*n= 78*) and transposases protein sequences (*n= 66*) as recently reported by Roy and co-workers (Roy et al., 2021). In this study, however, the inclusion of these protein sequences within the non-*intI*1 database is not of significance, as the goal was to confirm that analysed *intI*1 primer sets were unable to amplify sequences within this database via *in-silico* testing, thus confirming their specificity.

**Figure 1:**
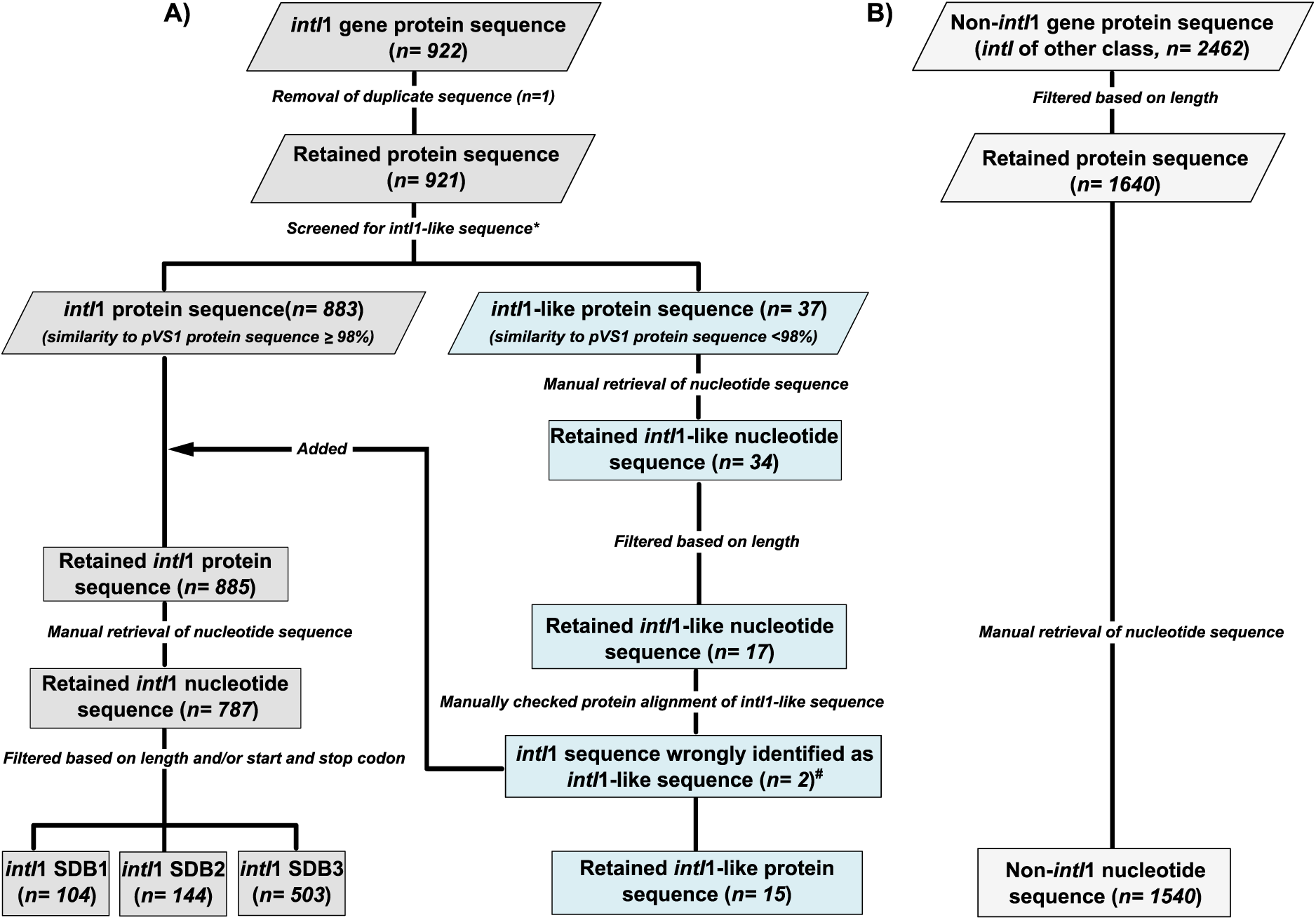
Workflow of integrase sub-databases construction for primer evaluation from 922 *intI*1 A) and 2462 non-*intI*1 protein sequences B) protein sequences. Duplicate *intI*1 protein sequence (*n=1*) was discarded. Retained protein sequences were compared to the reference *intI*1 protein of pVS1 plasmid (AAA25857.1) using NCBI BlastP to identify true *intI*1 sequences. *intI*1 protein sequences with a ≥ 98% identity to the pVS1 protein sequence were classified as *intI*1 sequence. Conversely, *intI*1 protein sequences with a <98% identity to the pVS1 protein sequence were classified as *intI*1-like sequences. Three *intI*1 sub-databases (SDB1, SDB2, SDB3) were finally constructed based on criteria specified in **Table 1** and were used to evaluate coverage of primers. *intI*1-Like (n= 15) and non-*intI*1 (*n=1540*) nucleotide sub-databases were used to evaluate specificity of primers. * Indicates removal of 1 protein sequence (CP006631.1) from the 921 non-duplicate *intI*1 protein sequence, due to no similarity score to the *intI*1 pVS1 protein sequence generated, as a result of low sequence similarity. ^#^ Indicates the two (WP_058137959.1 and WP_058135314.1) *intI*1 protein sequence incorrectly identified as *intI*1-like protein sequence by the low similarity score generated by NCBI following alignment to the pSV1 protein sequence due to these sequences being partial length sequences.

**Table 1:** Criterion for Construction of Integrase Sub-databases.

| Sub_databases<br>ID | Criteria |  |  | Number of sequences<br>within sub_database |
| --- | --- | --- | --- | --- |
|  | Nucleotide sequence<br>length (bp) | Beginning start<br>codons | Ending stop<br>codons |  |
| *intI1* SDB1 | $\geq 1000$ | ATG, TTG, GTG | TAA, TAG, TGA | 104 |
| *intI1* SDB2 | $\geq 900$ | ATG, TTG, GTG | TAA, TAG, TGA | 144 |
| *intI1* SDB3 | $\geq 600$ | N/A | N/A | 502 |
| *intI1*_like | $\geq 600$ | N/A | N/A | 16 |
| Non-*intI1* SDB | $>900$ | N/A | N/A | 1540 |

SDB= Sub-database

The *intI*1 database was curated by discarding duplicate protein sequences (*n=1*) from the 922 *intI*1 protein sequences (Figure 1). Retained *intI*1 protein sequences were then compared to the reference *intI*1 protein sequence of pVS1 plasmid (AAA25857.1) using NCBI BlastP, to ensure *intI*1 sequences used for *in-silico* assessment of primer and probe sequence coverage were indeed *intI*1 sequences as suggested by Roy et al., (2021). Further, protein sequences whose percentage identity to the reference pVS1 *intI*1 plasmid protein sequence was ≥ 98%, were characterised as *intI*1 sequence, while sequences whose protein similarity to the reference pVS1 *intI*1 plasmid protein sequence were <98% were categorised as *int*I1-like protein sequences (Roy et al., 2021). Moreover, protein sequence identified as *intI*1-like were manually checked to ensure percentage similarity score to the pVS1 protein sequence reported by NCBI was not due missing sequence caused by the alignment of partial sequence to a complete length sequence. As such, protein sequences (*n=2;* WP_058137959.1 and WP_058135314.1) incorrectly identified as *intI*1-like were added to the *intI*1 database (Figure 1).

Finally, three *intI*1 nucleotide sub-databases (SDB1, SDB2, and SDB3) were created for robust primer analysis based on the criterion specified in Table 1.

SDB1 (*n=104*) contained full length *intl*1 sequences ≥1000 bp, confirmed by the presence of a start and stop codon; SDB2 (*n=144*), contained full length *intl*1 sequences ≥ 900bp confirmed by the presence of a start and stop codon. Sequences within SDB1 are all present in SDB2. The final *intI*1 sub-database (SDB3, *n=503*) contained both complete and partial sequences (Figure 1; Table 1). All sequences within SDB1 and SDB2 were also present within SDB3. The *intI*1-like (<98% similarity to pVS1 on protein level) sub-database contained both complete and partial-length sequence (*n= 15*; Figure 1; Table 1). The-non *intl*1 database contained 1540 integrase sequences of other classes (Figure 1).

In parallel, the non-*intI*1 sequence sub-database was constructed from the 2462 *int*I of other class protein sequences by applying a ≥ 300 amino acid length thresholds (900bp nucleotide length) to filter out shorter-length protein sequence (Figure 1; Table 1). Retained protein IDs for the *intI*1, *intI*1-like and non-*intI*1 sequences were then used to manually obtain the nucleotide sequences from NCBI in Fasta format.

To summarise, *intI*1 sequences from this study was defined as *intI*1 protein sequences whose percentage identity shared a ≥ 98% similarity to pVS1 *intI*1 plasmid protein sequence (AAA25857.1), whilst *intI*1-like sequences was defined as *intI*1 protein sequences sharing a <98% similarity to pVS1 *intI*1 plasmid protein sequence (Figure 1).

#### 2.1.3 Primer evaluation

Published *intI*1 primers were analysed as primer pair (Table S.1), using Primer Prospector (Walters et al., 2011), to evaluate coverage and specificity against constructed integrase sub-databases (Figure 1). The analyze_primers.py function with the default settings on Primer Prospector was used to generate an alignment profile file for each primer against unaligned individual nucleotide sequence in each test sub-databases. For each primer alignment to a nucleotide sequence a weighted score (WS) was given.

**Overall WS was calculated as:**

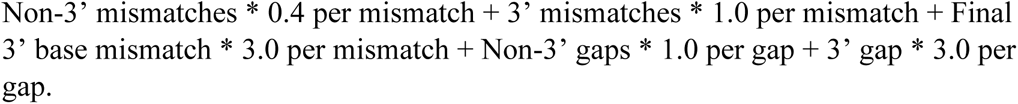

The first 5 bases of the primer and the target sequence were defined as the 3’end and thus, mismatches within these bases were termed 3’ mismatches. The remaining bases of the primer and the target sequence were defined as the non-3’ end. Therefore, mismatches within these non-3’ end bases were regarded as non-3’mismatches. Gaps in the alignment of the primer and the target sequence in the first 5 bases were termed 3’ gaps while gaps in the alignment for the remaining primer and template sequence were known as non-3’gaps. The lower the WS, the better the compatibility between the primer and target DNA sequence, a 0 score indicates perfect alignment. Primer Prospector will force a primer sequence to bind anywhere within the target sequence even if the primer binding site is unavailable to generate a WS for the primer. As such, the primer binding orientation of each analysed primer pair were checked for each sequence using R (R Core team 2022). In addition, the seqnir package(Gouy et al., 1984) in R was used to load the DNA/ protein sequences. The mean WS for the forward and reverse primer for the primer pairs with the correct-binding orientation was noted and a WS plot of each primer set generated using the “ggplot2” R package (Wickham, 2009). Detailed description of the primer evaluation can be found in the supplementary material 2.1.

In the case of primer pairs that incorporated a *Taq*Man hydrolysis probe, the primer-probe-binding-orientation (forward, probe and reverse) against each unaligned sequence was first verified, for each unaligned sequence by checking the hit positions of the forward, probe, and reverse primer sequence in R. Unaligned sequences with correct primer-probe orientation were subsequently retained and analysed in the manner as described in supplementary material 2.1.

#### 2.1.4 Design of a new *intI*1 primer set and *Taq*Man-minor-groove binder (*Taq*Man-MGB) probe

To improve *intI*1 sequence coverage and specificity for Q-PCR the *intI*1 primer set F3-R3, (Rosewarne et al., 2010, Table S.I) was modified to generate a new *intI*1 primer incorporating an MGB *Taq*Man probe set (*intI*1 DF-DR, Table S.I) following guidelines for primer-probe design outlined by McKew and Smith (McKew and Smith, 2015). An MGB probe of 15bp was designed using Primer Express software (Version 3.0.1; Applied Biosystems)^TM^. Detailed protocol of the MGB probe design can be found in the supplementary material 2.2. Primer and probe sequence were BLAST searched (BLASTN) to validate the sequence specificity. Specificity and coverage of the newly designed primer and probe set was tested as detailed above with Primer Prospector.

### 2.2 Validation of selected *intI*1 primers from in silico analysis on wastewater samples

#### 2.2.1 Optimisation of selected primer sets for Q-PCR

The amplicon produced from selected primers for laboratory validation were assessed *in-silico* first using sequences within SDB1 and then in the laboratory by end-point PCR. Selected *intI*1 primer sets that resulted in the correct size amplicon were further optimised for Q-PCR assays (Table 3). Q-PCR standard curves were constructed by amplifying a synthetic *intI*1 gene fragment (Integrated DNA Technologies) containing the primer binding site for all selected primers (Figure S.2). The insert fragment was amplified by PCR using T7 forward (5’-TAATACGACTCACTATAGGG-3’) and M13 reverse (5’-CAGGAAACAGCTATGAC-3’) primers. Reaction volume and condition in supplementary material 2.3. Resulting amplicons were purified, and size selected with the Agencourt AMPure XP beads (Beckman Coulter, Brea, CA, USA) per manufacturer’s recommendation, using a 1:1 ratio of beads volume to PCR product volume, and eluted in a 25μl volume nuclease-free water. Purified products were quantified fluorometrically using Qubit (Invitrogen, according to the manufactures recommendations) and gene copy number determined using EndMemo DNA copy number calculator (http://endmemo.com/bio/dnacopynum.php). The purified concentrated stock was subsequently diluted to 10^9^ copies/μl, followed by a five, 10-fold serial dilution (10^7^-10^3^ copies/μl) for amplification by Q-PCR. Standard curve was obtained by plotting the average of each triplicate threshold cycle (Cq) against the log10 of standard concentration (copies/μl). Standard curve descriptors including efficiency, slope, y-intercept and R^2^ are reported (Smith and Osborn, 2009).

**Table 2:** Selected Sample Timepoint for Each Septic Tank Investigated.

| Reactor type | Reactor ID | April 2018 | May 2018 | June 2018 | Nov 2018 | March 2019 | June 2019 | July 2019 | August 2019 | Sept 2019 |
| --- | --- | --- | --- | --- | --- | --- | --- | --- | --- | --- |
| Solar septic tank (SST) | SST-01 | EFF/SLG | EFF/SLG | EFF/SLG |  |  |  |  |  |  |
|  | SST-07 | EFF/SLG |  |  | EFF/SLG | EFF/SLG |  |  |  |  |
| Conventional septic tank (CST) | CST-P3 |  |  |  |  |  | INF/EFF/SLG | INF/EFF/SLG | INF/EFF/SLG |  |
|  | CST-J6 |  |  |  |  |  | EFF/SLG | EFF/SLG | EFF/SLG |  |
|  | CST-HC |  |  |  |  |  |  |  | EFF/SLG | EFF/SLG |

INF Influent; EFF Effluent; SLG Sludge. SST Solar septic tank; CST Conventional septic tank
† Excluded from *intI1* Q-PCR quantification due to insufficient sample volume but was included in MiSeq amplicon sequencing

**Table 3:**
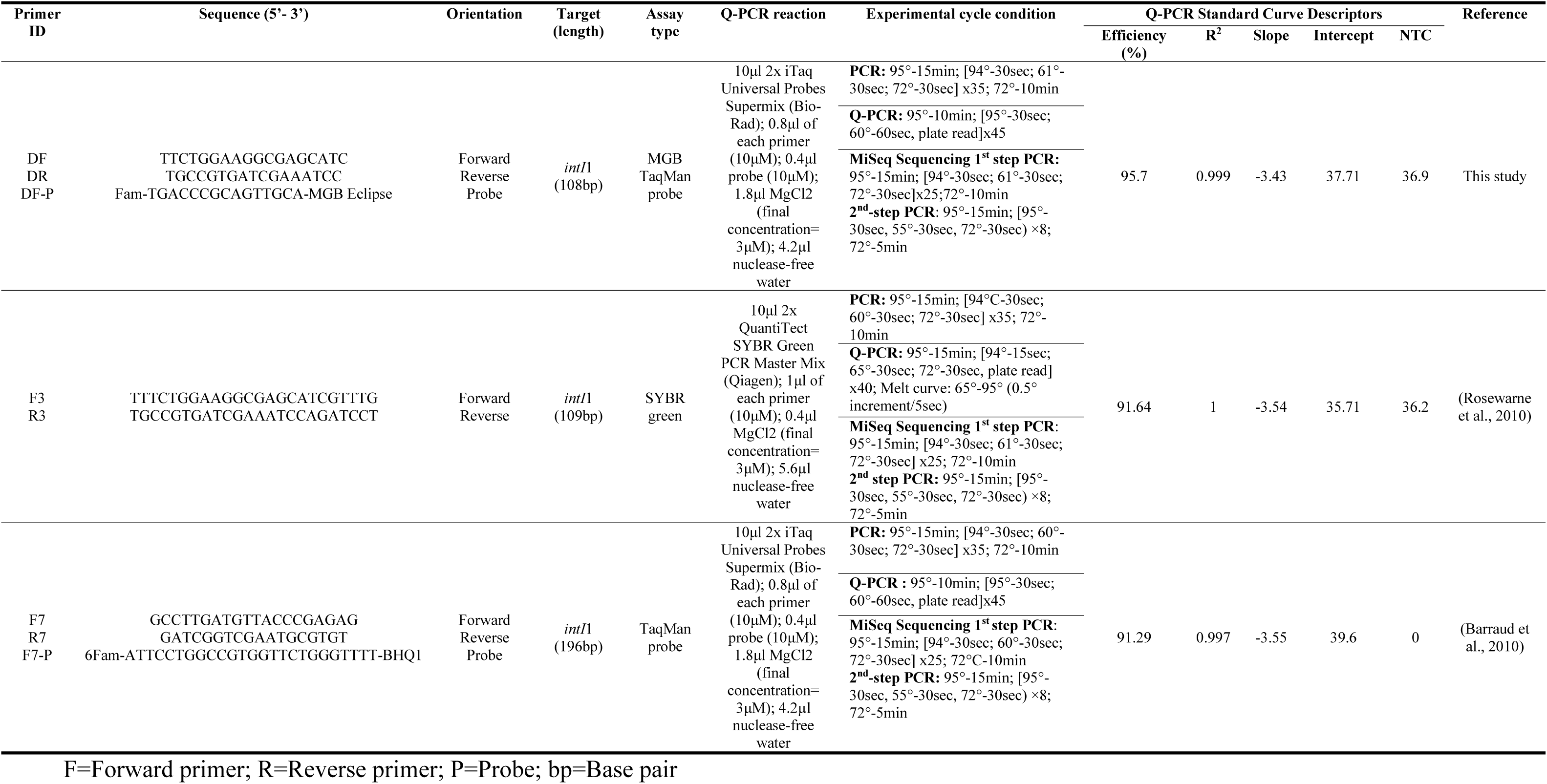
Primers and probe sets selected and optimised for Q-PCR to quantify *intI*1 gene copies from wastewater.

### 2.3 Application of selected *intI*1 primers from SST and CST wastewater samples

#### 2.3.1 Solar and Conventional tank sampling

Two household solar septic tank (SST; SST01 and SST07) units and three conventional septic tank (CST; two household tanks and one healthcare tank) units, operational within the Pathum Thani province and Samut Prakan province, Thailand, were sampled between April 2018 to September 2019 (Table 2).

The SST and the household CST units (CST-P3 and CST-J6) have a 1000L total working capacity, whilst the healthcare CST units (CST-HC2; herein referred to as CST-HC) has a 2000L total working capacity each. Each tank was buried to approximately 1.5 metres below ground level; with the tank surface (lid) at ground level, and so exposed to atmospheric temperatures (Connelly et al., 2019). The sampling approach used is described in **supplementary material 2.4** is outlined elsewhere (Connelly et al., 2019). 100 ml of effluent and 40 ml sludge was sampled from the SST and CSTJ7 and CST-HC2, while 100ml of influent was also collected from CST-P3. 40 ml of sludge was sampled from each reactor. All samples were pelleted for DNA extraction. The months for sampling the SST were selected based on the highest recorded internal temperature of the 12-month sampling campaign conducted.

#### 2.3.2 DNA extraction

From each sample, DNA extraction was performed with the DNeasy PowerSoil Kit (Qiagen), in accordance with the manufacturer’s instructions. Integrity of extracted genomic DNA was assessed via agarose gel electrophoresis and DNA concentration quantified fluorometrically using the Qubit (Invitrogen) according to manufactures instructions.

#### 2.3.3 Q-PCR quantification of *intI*1 gene from wastewater

*intI*1 genes were quantified from septic tank wastewater samples from Thailand (Table 2) using optimised Q-PCR conditions for the three selected *intI*1 primer pairs (DF-DR, F3-R3 and F7-R7). For each primer set, Q-PCR amplification was carried out in a 20μl volume reaction using 2µl (1:50 diluted) template DNA. Reaction volume, conditions, primer sequences and probe type for the three selected optimal *intI*1 primer-pairs are detailed in Table 3. Triplicate/ duplicate no template control (NTC) was included for each primer set. Reactions were performed on the Bio-Rad CFX96 Touch Real-Time PCR Detection System and analysed with the Bio-Rad CFX Manager 3.1 software. Melt curve analysis was performed, for the SYBR Green assay, from 65°C to 95°C with 0.5°C increments every 5 secs, and a single peak confirmed to ensure assay specificity.

Statistical analyses were performed in R (R Development Core Team, 2008). Two-way analysis of variance (ANOVA) followed by a Turkey HSD post hoc test, was employed to compare *intI*1 gene abundance for each of the sample types (influent, sludge, and effluent) and reactor type (CST and SST) for each primer set. Finally, a Pearson correlation analysis was applied to calculate the relationship between the abundance of *intI*1 detected between each primer set.

#### 2.3.4 MiSeq Amplicon sequencing

The specificity of the selected *intI*1 primer sets (DF-DR, F3-R3 and F7-R7) used to quantify *intI*1 gene from septic tank sludge and wastewater, were confirmed by Illumina MiSeq amplicon sequencing of the *intI*1 gene from 31 wastewater samples (Table 2) using the optimised endpoint PCR conditions outlined in Table 3. A two-step PCR was performed to barcode samples as detailed previously (Bourlat et al., 2016; Cholet et al., 2019). Detailed description of the method is provided in the supplementary material 2.5.

#### 2.3.5 Bioinformatics

Primer sequences were used to extract the *intI*1 gene from the resulting reads, particularly for shorter primer pairs, using the Cutadapt algorithm (Martin, 2011). Abundance tables were then generated by constructing amplicon sequencing variants (ASVs) using the Qiime2 pipeline and the DADA2 algorithm (Bolyen et al., 2019) with details given at https://github.com/umerijaz/tutorials/blob/master/qiime2_tutorial.md. Constructed ASVs was blast searched on NCBI and closest hit sequences retrieved for each ASV. Phylogenetic distance between sequences was investigated. First, a multiple sequence alignment of ASV sequences, retrieved NCBI sequences, complete length *intI*1 and *int*I1-like and an *intI*3 (class three integrase gene; nucleotide ID: AY219651.1) sequence was done using MAFFT (Katoh et al., 2002) for each primer set. Aligned sequences were visualised in BioEdit (version 7.0.5.3) (Hall, 1999) and trimmed to retain only aligned region without gaps. Phylogenetic trees were constructed using maximum likelihood approach with a generalised timer-reversible substitution model implement in RAxML version 8 (Stamatakis, 2014). Consensus trees were calculated after 1000 bootstrapping permutations.

Phylogenetic tree of the trimmed and aligned sequence, for each primer pair, was constructed with RAxML (Price et al., 2009). A heat tree of the constructed ASVs, after log2 transformation of ASVs abundance per sample for each primer set, was mapped to analysed samples, coloured, and visualised using the ggtree package (Yu et al., 2017). Tip of tree was coloured based on sequence isolation source.

### 2.4 Validation of selected primers to quantify *intI*1 mRNA transcripts from environmental samples

#### 2.4.1 Sample collection, filtration, and DNA/RNA co-extraction

As the septic tank wastewater samples were previously collected and only DNA extracted and stored at −80°C, they were not suitable for RNA analysis (Cholet et al., 2019). Therefore, we tested the suitability of the optimised primers sets to detect *intI*1 mRNA using freshly collected environmental samples of river water collected from the Kevin River, Glasgow (UK), to determine if *intl*1 mRNA transcripts could be quantified in receiving water bodies. 3L of surface water was collected in March and April 2022 and filtered through a sterile glass microfibre filter (FisherBrand MF200; retention 1.2μm) and onto a 0.22μm Sterivex filter. Filters were immediately extracted from or frozen at −80°c for later use.

DNA-RNA co-extraction was carried out according to protocol previously described(Griffiths et al., 2000; Tatti et al., 2016; Cholet et al., 2019), with a minor modification to the bead-beating time (45 sec) as outlined by Lim et al., (2016). Detailed description of this method is outlined in supplementary material 2.6. Briefly, RNA was prepared from the DNA-RNA co-extract by DNase treating with Turbo DNase Kit (Ambion) in accordance with the manufacturer’s recommendation, with modification to the incubation time and volume of DNase added as previously described (Cholet et al., 2019), 1μl DNase volume was added and incubated at 37°c for 1 hour, followed by further addition of 1μl DNase and a re-incubation at 37°c for another hour. Detailed protocol can be found in the supplementary material 2.7.

#### 2.4.2 RT-Q-PCR quantification of *intI*1 genes and transcripts from river water

Q-PCR DNA standard curve were constructed as above (**see above section 2.2.1**). For each primer set*, intI*1 cDNA and DNA Q-PCR amplification was carried out in a 20μl volume reaction using 2µl (1:2 and/ 1:5 diluted) template DNA/ cDNA. In addition, two priming strategies, Gene specific (GS) and/ or Random (RH) priming were used to reverse transcribe *intI*1 mRNA to cDNA. Detail of this approach provided in supplementary material 2.7. Q-PCR reaction volume, conditions, primer sequences and probe type for the three selected optimal *intI*1 primer-pairs are the same as specified in Table 3. Reactions were performed on the Bio-Rad CFX96 Touch Real-Time PCR Detection System and analysed with the Bio-Rad CFX Manager 3.1 software. Melt curve analysis was performed, for the SYBR Green assay, from 65°C to 95°C with 0.5°C increments every 5 secs, and a single peak confirmed to ensure assay specificity. Standard curve descriptors including efficiency, slope, y-intercept and R^2^ are reported (Table 4).

**Table 4:**
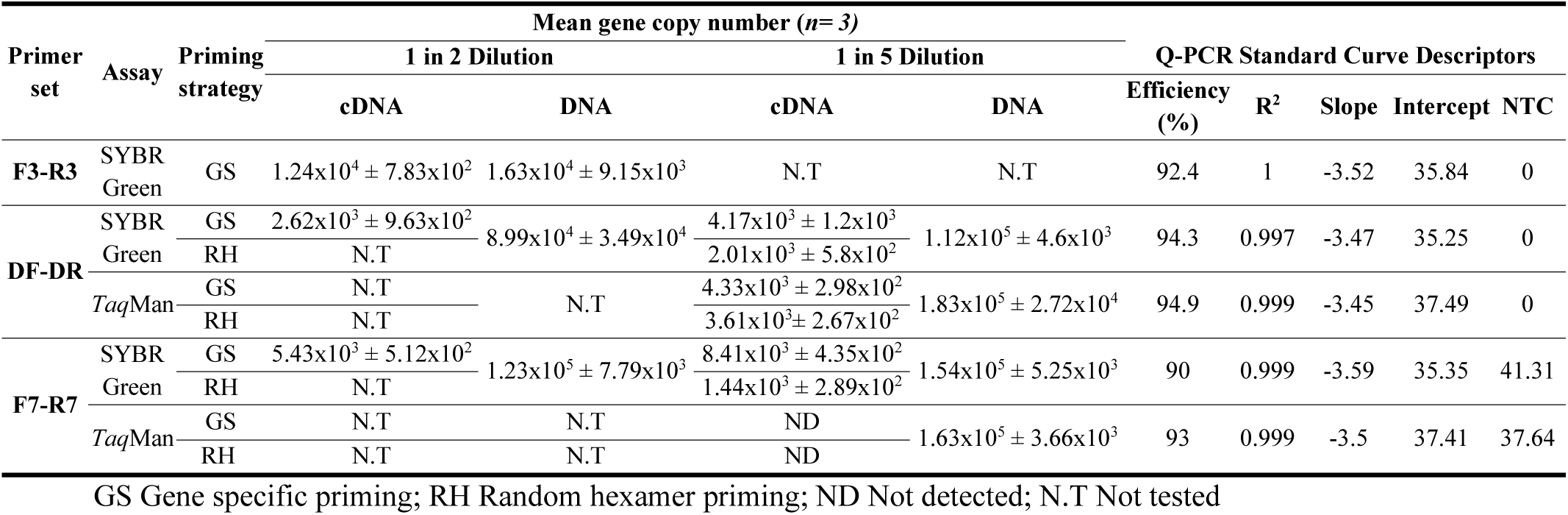
*intI1* mRNA transcripts copies/ng DNA.

## 3.0 Results

### 3.1 *intI*1 primer evaluation

#### 3.1.1 Evaluation of primers for coverage

In total 64 different *intl*1 primer sets, including 4 *Taq*Man primer-probe sets were retrieved from the systematic review (Supplementary Table S.1). In addition, the primer and probe set designed in this study was included in the analysis, resulting in 65 primers evaluated (Table S.1). Primers were initially aligned against the reference *Pseudomonas aeruginosa* plasmid pVS1 nucleotide sequence (M73819.1) (Supplementary Figure S.1) and renamed for ease of identification (Supplementary Table S.1). Next, primers were aligned against the SDB1 (full length *intl*1 database) to ensure binding sites were present (in forward or reserve orientation) and that the expected amplicon size would be generated. From this, 10 primer sets were discarded, which included two sets (F61-R61 and F64-R64) that were not *intI*1 primers (Supplementary Table S.2). The F61-R61 primer set targeted *aadA1a* aminoglycoside adenylyl transferases gene (Sandvang et al., 1997; Guerra et al., 2001), while the F64-R64 primer pair targeted the class two integron-integrase gene (*intI*2) (Gündoǧdu et al., 2011). In addition, these primer sets (F61-R61 and F64-R64) aligned poorly to the reference *Pseudomonas aeruginosa* pVS1 *intI*1 nucleotide sequence (data not shown) and had no hit (High WS) with complete length *intI*1 sequences within SDB1 (Supplementary Table S.2).

Of the remaining 55 primer pairs, when aligned to SDB1 (complete length sub-database *n=104*) ≥ 97-100% of sequences had the correct primer binding orientation (Supplementary Table S.3).In addition, the majority of these primers (49 sets; 89%) amplified 69-100% of sequences in the correct primer binding orientation with 0 mismatch (Supplementary Figures S.3A, S.3B; Supplementary Table S.3). One primer (F40-R40) performed poorly, amplifying only 0.9% (*n=1* amplicon) sequences with correct primer binding orientation at 0 mismatch (Supplementary Figure S.3; Supplementary Table S.3), and was removed from further consideration.

Primer coverage was then tested against the other *intI*1 complete length and partial length sub-databases, SDB2 (*n=144*) (Supplementary Figures S.3C, S.3D) and SDB3 (*n=503*) (Supplementary Figure S.4).respectively. Here, the number of sequences with the correct primer binding orientation declined (SDB2: 79-100%; SDB3: 41%-100% of the sequences had the correct primer binding orientation), as did the number of amplicons amplified with a 0 mismatch (SDB2: 50-99%; SDB3: 16%-99%) for the correct primer binding orientation sequences (Supplementary Figures S.3, S.4; Supplementary Table S.3).

Five (9%) primer sets produced no amplicon at a WS of 0 across the three *intI*1 sub-databases (Supplementary Figure S.3; S.4; Supplementary Tables S.3) and were removed from further consideration. This included one set (F29-R29) which performed optimally when allowing a single mismatch at the 5’ end (WS=0.4) (Supplementary Table S.4), but as there were several primers with better coverage at a WS of 0, this primer set was removed. In summary, a further 6 primer sets (F9-R9, F29-R29, F40-R40, F53-R53, F56-R56, F62-R62) were discarded from the primer coverage analysis.

Of the 49 amplicon-producing primers at 0 WS retained, 10 primer sets (DF-DR, F1-R1, F3-R3, F7-R7, F13-R13, F16-R16, F31-R31, F35-R35, F57-R57 and F60-R60) consistently had a high number (99-100%) of sequences with the correct primer-binding orientation and amplified ≥97% amplicons within the complete length sub-databases (SDB1: *n=104* and SDB2 *n=144*). Moreover, these primers consistently had a low mean WS for the forward and reverse primer within each pair (Supplementary Table S.3). As such, these 10 primers were considered the best performing *intI*1 primer sets.

Five primer sets analysed in this study incorporated a *Taq*Man probe (Supplementary Table S.5), two of which (DF-DR and F7-R7 sets) were among the best performing primer set. Of these, the DF-P-DR primer-probe set, designed in this study, consistently produced the highest number of amplicons at 0 WS across the three *intI*1 sub-databases with 102 (98%), 142 (99%) and 494 (99%) of sequences amplified within SDB1, SDB2 and SDB3 respectively (Supplementary Table S.5). In addition, allowing for a single non 3’ mismatch (WS=0.4) between primer and probe, resulted in all sequences with the correct primer-binding orientation to be amplified across the three *intI*1 sub-databases (Supplementary Table S.5).

Conversely, primer-probe set F46-P-R46 performed the worst, of five primer-probe combinations assessed, with only 34 (33%), 41 (32%) and 88 (23%) of sequences with the correct primer-binding-orientation amplified at 0 WS within SDB1, SDB2 and SDB3 respectively. Nonetheless, allowing for an increased WS of 1 (i.e., mismatch caused by either a single 3’ end mismatch, a non-3’end gap or two non-3’ mismatches) resulted in significant increase in the number of amplicons amplified across *intI*1 sub-databases (SDB1: *n=95* (93%), SDB2: *n=121* (93%), SDB3: 360(95%)) (Supplementary Table S.5).

The F7-P-R7 primer-probe set (commonly used *Taq*Man assay in *intI*1 gene study), F10-P-R10 and F38-P-R38 primer-probe sets showed similar coverage to each other, but lower than DF-P-DR set, with 92 (88%), 91 (88%) and 92 (88%) of amplicons amplified at 0 WS within SDB1 respectively (Supplementary Table S.5). However, the F7-P-R7 primer-probe set amplified the highest number of amplicons at 0 WS (or second highest after the DF-DR set) for the correct primer binding sequences across the other two *intI*1 sub-databases (SDB2: *n=131* (91%), SDB3: *n=454*(96%)) among the three primer sets (Supplementary Table S.5). As such, the DF-P-DR and F7-P-R7 primer-probe sets were put forward as the top performing primer-probe set.

#### 3.1.2 Evaluation of primers for specificity

The primer sets were tested for specificity against the *intI*1-like (*n= 15*) and non-*intI*1 (*n= 1540*) sub-databases respectively (Supplementary Figure S.5; Supplementary Table S.3). Here, the aim was for the primers to amplify the least amount of non-target sequence reflected by a higher forward and reverse primer WS for sequences where primers bind in the correct orientation. The 10 best performing primer sets identified above were focused on.

For the best performing primer sets, the number of sequences with correct primer-binding orientation ranged from 67-100% and 41-65% for the *intI*1-like and non-*intI*1 sub-databases respectively with 57-80% and 0% of these correct primer-binding orientation sequences amplified at 0 mismatch in *intI*1-like and non-*intI*1 sub-databases respectively (Supplementary Figure S.5, Supplementary Table S.3).

Of these best performing sets, the F16-R16 primer set amplified the highest number of *intI*1-like amplicons (*n=11*, 79%) at 0 mismatch and was removed, while the primer pair F57-R57 amplified the lowest number of *intI*1-like sequence (*n=7*) (Supplementary Figure S.5A, Supplementary Table S.3). The incorporation of a *Taq*Man probe generally improved primer specificity, however two of the primer sets which incorporated a *Taq*Man probe (DF-P-DR and F7-P-R7) both amplified *intI*1-like sequence. The number of *intI*1-like amplicons amplified by the DF-P-DR (*n=9*) and F7-P-R7 (*n=8*) primer-probe sets at a 0 WS were similar. Of note, whilst the 10 best performing was focused on, the other primer sets analysed (**see section 3.1.1**) also amplified *intI*1-like sequences (Supplementary Figure S.5A; Supplementary Table S.3).

Next, the nine remaining primer sets were tested against the non-*intI*1 sequences (Supplementary Figures S.5B, S.5C). None of the primers amplified non-*intI*1 sequence at a 0 mismatch. In general, primers only produced amplicons from the non-*intI*1 database with very high weighted scores (sum of forward and reverse primer mean WS ranged: 8.39-11.6) (Supplementary Figures S.5B, S.5C; Supplementary Table S.3). However, the primer pairs F1-R1 (WS: 2) and F13-R13 (WS: 3.2) performed worst, having the lowest WS required to amplify at least one non-*intI*1 target. As such, were removed from further analysis, leaving seven sets (DF-DR, F3-R3, F7-R7, F31-R31, F35-R35, F57-R57 and F60-R60) to be considered the best overall performing *intI*1 primer sets in terms of coverage and specificity.

#### 3.1.3 Recommendation of optimal primer sets for *in situ* laboratory validation and *in-silico* amplicon size distribution

From the initial 65 primer sets, seven (DF-DR, F3-R3, F7-R7, F31-R31, F35-R35, F57-R57 and F60-R60) were identified that had high coverage in our *intl*1 database, but low-specificity to the non-*intI*1 database, indicating they are good primer sets targeting a broad range of *intl*1 targets, while discriminating against non-*intI*1 sequences. Two published sets (F3-R3 and F7-R7) were selected (Supplementary Table S.3), as they not only had the highest WS required to amplify non-*intI*1 target (i.e., needed the highest number of mismatches to target the sequence; F3-R3-WS: 5.2 (2 non-*intI*1 amplicons), F7-R7-WS 5.2: (6 non-*intI*1 amplicons)) but also had short amplicons (100-200bp), making them ideal for both Q-PCR and high-throughput amplicon sequencing. In addition, each of these selected primer sets targeted a different region of the *intl*1 gene and were commonly used within the literature (Barraud et al., 2010; Rosewarne et al., 2010; Stalder et al., 2014; Paiva et al., 2015; Johnson et al., 2016; McKinney et al., 2018). F7-R7 incorporated a *Taq*Man probe. The primer and probe set, DF-P-DR, designed in this study, was also included resulting in three *intI*1 primer sets selected for laboratory validation.

### 3.2 Application of selected *intI*1 primers on septic wastewater samples

#### 3.2.1 Q-PCR quantification of *intI*1 gene from Thai Septic Tanks wastewater

The three selected and optimised *intI*1 primer sets and probes (DF-DR, F3-R3 and F7-R7; Table 3) were used to quantify *intI*1 gene abundance across 30 septic tank wastewater samples (influent, sludge and effluent) from CST-household, CST-healthcare and SST-household reactors (Table 2).

Each of the standard curves from all three primer sets had high efficiencies which ranged from 91.29 to 95.7%, y-intercepts of 35.71 to 39.6, slope of −3.43 to −3.55 and a No Template Control Ct from undetected to 36.9 (Table 3).

Of note, between primers sets, there was no difference in *intI*1 gene abundance for the same sample type (influent, sludge, effluent) (p-value >0.05, Figure 2; Supplementary Table S.6), and Pearson correlation coefficient analysis indicated that the *intI*1 gene copy number amplified by each of the primers were highly correlated (r=0.982 (p-value <0.001), r=0.99 (p-value <0.001), and r=0.993 (p-value <0.001) for DF-DR and F7-R7, DF-DR and F3-R3, and F3-R3 and F7-R7 primer sets respectively). As such, each primer set resulted in the same overall pattern of *intI*1 gene abundance, with higher *intI*1 gene copies/ng DNA observed in the **influent** (DF-DR: 4.22×10^4^±SD3.57×10^4^; F3-R3: 3.66×10^4^±SD3.06×10^4^; F7-R7: 4.89×10^4^±SD4.29×10^4^ copies/ng DNA) > **effluent** (DF-DR: 3.33×10^4^±SD2.30×10^4^; F3-R3: 2.88×10^4^±SD1.93×10^4^; F7-R7: 3.53×10^4^±SD2.39×10^4^ copies/ng DNA) > **sludge** (DF-DR: 8.81×10^3^±SD3.94×10^3^; F3-R3: 7.72×10^3^± SD 2.65×10^3^; F7-R7: 8.24×10^3^±SD 3.07×10^3^ copies/ng DNA) (Figure 2).

**Figure 2:**
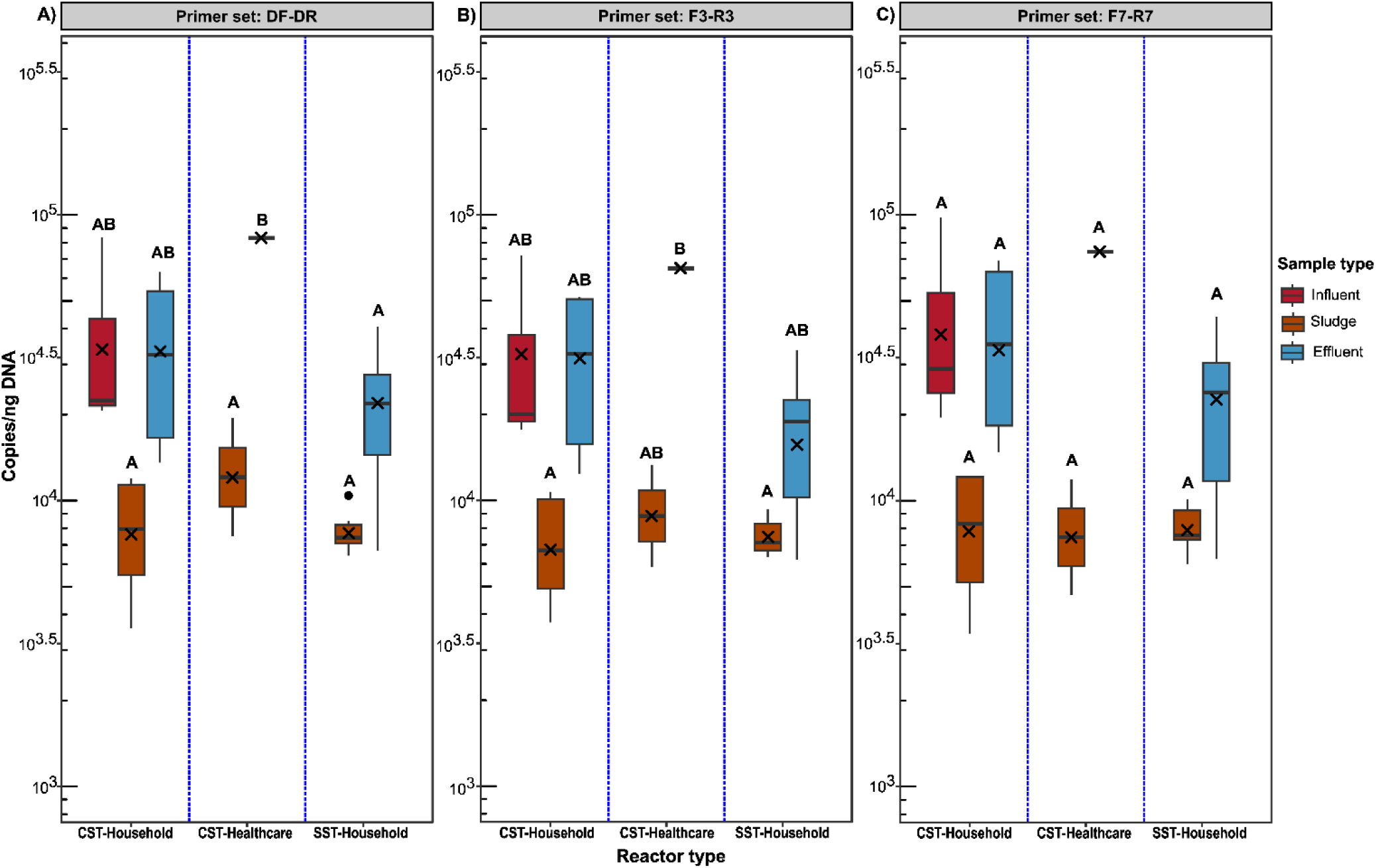
Impact of primer choice on the quantification of *intI*1 gene copies from CST-Household, CST-Healthcare and SST-Household septic tank wastewater reactors, and three wastewater sample types (influent, effluent, sludge). Results of the two-way ANOVA analysis showing statistically significant difference in *intI*1 gene copies quantified between reactor types and sample types. For each primer set, boxplot sharing the same letter indicates no statistically significant difference at p-value >0.05, while boxplot with different letters indicates statistically significant difference at p-value <0.05. Statistically significant difference in *intI*1 gene abundance between primer sets for the same sample was not observed (p-value >0.05; see supplementary Table S.VI). X icon indicate mean *intI*1 copy number/ng DNA. Black dot indicates data outlier.

Although a similar overall pattern of *intI*1 gene abundance was observed, the F7-R7 primer set was the only primer set that reported no statistical difference (p-value > 0.05) in gene abundance between samples (influent, sludge, effluent) and reactors (CST-Household, CST-Healthcare and SST-Household), while only DF-DR primer set showed significantly higher *intl*1 gene abundances in the effluent of the CST healthcare than all other samples (Figure 2).

SST-household units incorporated an internal pasteurisation effect and were therefore expected to have lower *intl*1 gene abundance in the effluent. *intI*1 gene abundance per ng of DNA were lower in effluent than in both the CST-household and CST-healthcare tanks for all three primer sets (Figure 2). However, these differences were only statistically significant for DF-DR primer se (p-value= 0.005) between SST-household and CST-healthcare effluent (Figure 2). Nonetheless, the lower *intI*1 gene abundance quantified in the solar septic tank (SST-household) effluent, albeit only statistically lower in the CST-healthcare for the DF-DR primer set.

Although, *intI*1 gene copies in the sludge of the reactors where lower than the effluent, these *intI*1 gene copies were still high, albeit that the gene abundance between the three reactors was marginally different but not statistically significant (p-value >0.05), regardless of the primer set (Figure 2). However, depending on the primer set used, the reactor with the higher abundance and thus likely to contribute most to the environment changed.

The DF-DR primer set reported the CST-healthcare (1.35×10^4^±SD8.45×10^3^ copies/ng DNA) to be the higher contributor of *intI*1 gene to the environment via sludge and the SST-household unit (7.86×103±SD1.43×10^3^ copies/ng DNA) to be the lowest of the three reactors (Figure 2A), while primer set F3-R3 also indicated the CST-healthcare sludge as the higher contributor (9.58×10^3^±SD5.27×10^3^ copies/ng DNA) of CL1-intgeron to the environment via sludge, but reported the CST-household (7.30×103±SD3.11×10^3^ copies/ng DNA) as the least contributor (Figure 2B). Finally, the F7-R7 primer set revealed the CST-household unit sludge (8.42×10^3^±SD4.10×10^3^ copies/ng DNA) to be the greater contributor of *intI*1 to the environment and the SST-household (8.06×10^3^±SD1.59×10^3^ copies/ng DNA) as the least contributor (Figure 2C). As such, the SST-household in general had the lowest *intl*1 gene abundances in sludge when primer sets DF-DR and F7-R7 was used, but not when the F7-R7 primer set is used (Figure 2).

Influent samples were only accessible from the CST units with *intI*1 gene abundance higher in the **influent** (DF-DR: 4.22×10^4^ ±SD3.57×10^4^; F3-R3: 3.66×10^4^±SD3.06×10^4^; F7-R7: 4.89×10^4^±SD4.29×10^4^; copies/ng DNA) than **effluent** (DF-DR: 2.43×10^4^±SD1.76×10^4^; F3-R3: 2.52×10^4^± SD2.06×10^4^; F7-R7: 2.95×10^4^±SD2.34×10^4^ copies/ng DNA), indicating a removal efficiency ([influent-effluent/influent]*100) of 42.33%, 31.21%, 39.63%, for the DF-DR, F3-R3 and F7-R7 primer sets respectively (Figure 2).

In summary, primer sets used did not change the overall pattern of *intl*1 gene abundances nor did it result in statistical difference (p-value >0.05) in *intl*1 gene abundance for the same sample type (influent, sludge, effluent) quantified with the different primers. However, comparing samples within the same primer set did sometimes result in statistical differences between samples., which could alter interpretation of the risk of *intl*1 gene abundances, and in turn AMR pollution to the environment.

#### 3.2.2 MiSeq amplicon sequencing

MiSeq amplicon sequencing was undertaken on all septic tank samples (*n=31*; Table 2), to confirm the specificity of the selected *intI*1 primers (Table 3) and assess the diversity of the *intl*1 amplicons retrieved from the septic tanks. Overall, the number of unique ASV generated by each primer set was low, with 3 ASVs for DF-DR; 4 ASVs for F3-R3 and 11 ASVs for F7-R7 primer set (Supplementary Table S.7).

A phylogenetic tree was constructed with the highly conserved *intl*1 sequences from a range of environmental and clinical samples. In addition, *intl*1-like sequences (*n=4*) were added to the tree to determine if the primer sets could distinguish between these and *intl*1 sequences (Figure 3). All *intl*1 and *intl*1-like sequence, including the ASVs found here showed high sequence similarity to each other, likely owing to the short amplicon region designed over conserved regions (Figure 3). *intl*1-like sequences clustered among the *intl*1 and ASVs for all three primer sets, indicating that the primer sets could not differentiate between both variants.

**Figure 3:**
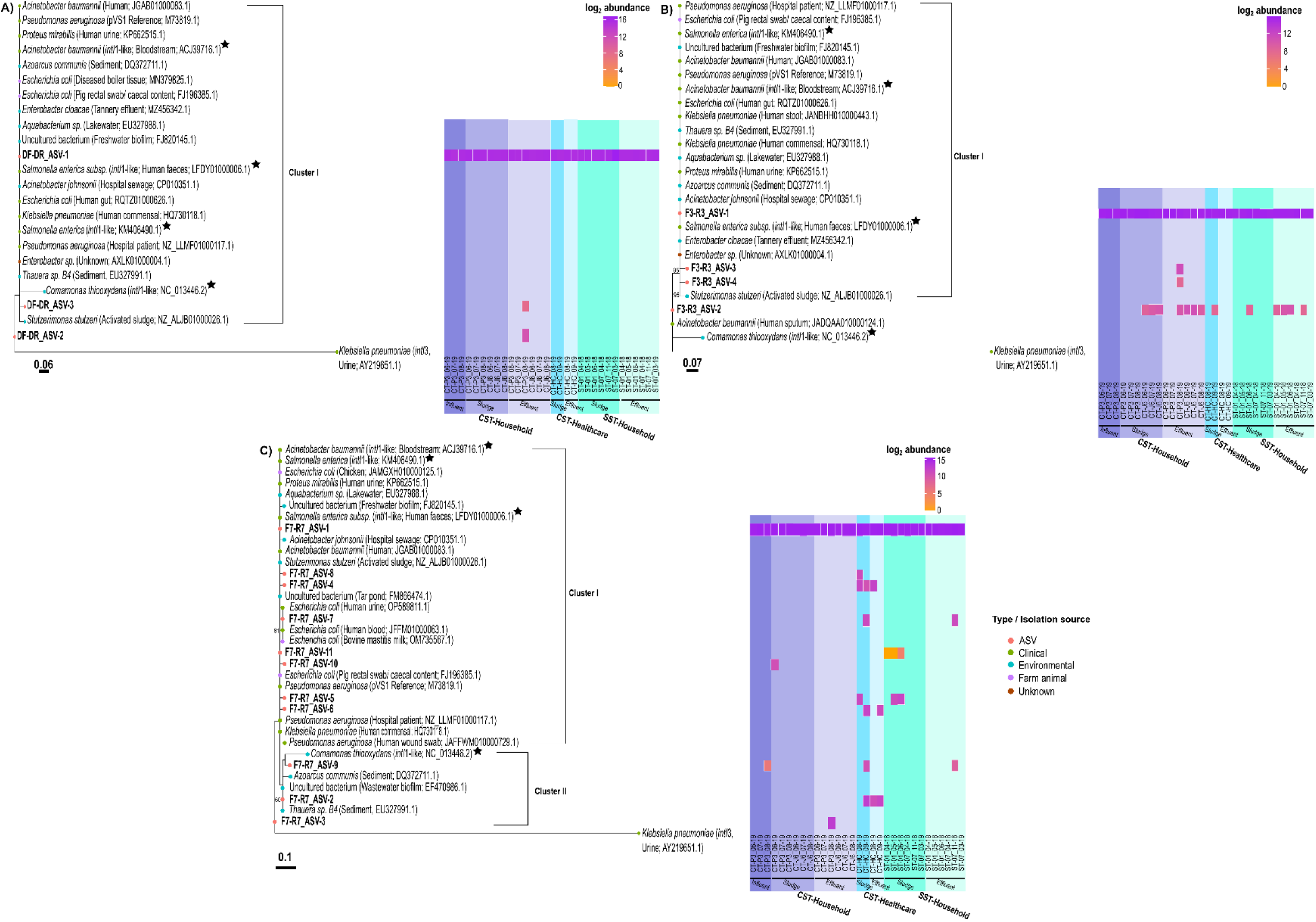
Detected ASVs abundance in Thai septic tank wastewaters (SST-Household, CST-Healthcare and SST-Household) by the DF-DR (A), F3-R3 (B) and F7-R7 (C) *intI*1 primer sets. Generated ASVs coupled with known and unknown *intI*1 within SDB1 (*n=104*), best hit NCBI sequences, and *intI*1-like sequences were aligned with Mafft, trimmed to only aligned region with no gaps, and phylogenetic tree constructed using the RAxML with 1000 bootstrap permutations. Number at node represents bootstrap value > 50% (from 1000 permutations). Bootstrap value at node <50 are not shown. The class 3 integron-integrase (*intI*3) gene (nucleotide ID: AY219651.1), which on protein level, shared a 60.74% similarity to the pVS1 protein sequence (AAA25857.1) was used as the outgroup. Colour of tree tips indicates isolate source of sequence/ ASVs generated by primer set. Heatmap shows log2 fold abundance (mean number of ASVs-DF-DR:5.1955×10^4^, F3-R3: 4.6602 x10^4^ and F7-R7: 3.6684 x10^4^; Table S.7) of detected ASVs within each wastewater sample. CTP3 and CTJ6 samples originated from two independent CST-Household reactor. CT-HC sample was from a CST-Healthcare tank. ST01 and ST07 are two independent SST-Household units. The sampling month and year is indicated by the format month_year (i.e., 06_19= June 2019). CST, Conventional septic tank; SST, Solar septic tank

All three primer sets amplified an abundant ASV-1 phylotype as the dominant *intI*1 sequence present in all septic tank types and sample types. This ASV was highly similar to *intI*1 and *intl*1-like sequences found in a range of environmental samples including freshwater biofilm, tannery effluent, hospital sewage and activated sludge (Figure 3).

For primer set DF-DR, a single cluster of *intl*1 sequences was present, albeit not supported by a bootstrap value, which also contained a second ASV (ASV-3) only present in the CST-household effluent (CST-P3_08-19). A third ASV (ASV-2), again only detected in the CST-household effluent (CST-P3_08-19), clustered outside the main group, highly similar to *intI*1 sequence from activated sludge, although with a low bootstrap value (Figure 3A).

For primer set F3-R3, a single cluster was observed, supported by a 95% bootstrap value containing ASV1, 3 and 4. These clustered with unknown and known *intl*1 from Tannery effluent and activated sludge as well as known *intI*1-like sequences of clinical origin (Figure 3). While ASV-1 was present in all samples, ASV3 and 4 were only detected in the CST-household effluent (CST-P3_08-19). Outside of this cluster was ASV2, highly similar to *intl*1 from *Acinetobacter baumannii*, a clinical pathogenic bacterium. ASV2 was present in both the CST-household and SST-household tanks sludge and effluent but only present in one CST-healthcare sludge sample (CT-HC_09-19) (Figure 3B). As primer sets DF-DR and F3-R3 targeted the same region of the *intI*1 (Figure S.1), the ASVs generated by each primer set (DF-DR and F3-R3-ASV1; DF-DR-ASV2 and F3-R3-ASV3; DF-DR-ASV3 and F3-R3-ASV4) had 100% sequence similarity to each other but only ASV1 from each primer set showed a 100% sequence similarity when aligned against the full length *intl*1 nucleotide sequences (pVS1, M73819.1).In addition, F3-R3-AV2 did not align to the full length *intl*1 with a 100% similarity.

Primer set F7-R7 detected 11 ASVs separated in two clusters. Within cluster I, ASV-1 present in all samples, clustered with ASV 8, 4, 7, 11, 10, 5 and 6 and was detected in CST-household (sludge), CST-healthcare (sludge and effluent) and SST-household (sludge and effluent) reactors. It clustered with known *intl*1 from sources such as hospital sewage and Tar-Pond, but also *intl*1-like sequences. Clustering was not supported by a high bootstrap value. Within cluster II, ASV-9 and 2 were detected in CST-household influent, CST-healthcare sludge and effluent and SST-household effluent samples and clustered with unknown and known *intI*1 sequence, as well as *intI*1-like sequence, found in environmental source such as sediment and wastewater biofilm. However, clustering was not supported by a high bootstrap value (<50%) (Figure 3). A final ASV (ASV-3), again only detected in the CST-household effluent (CST-P3_08-19), clustered outside the main group, but was not supported by a high bootstrap value (<50%) (Figure 3C).

In summary, *intl*1 diversity showed all samples to be dominated by a single ASV-1. It was highly similar to *intl-1* from clinical and environmental samples, however, *intl*1-like samples also clustered with it. CST-Household had the highest richness with primer set DF-DR and F3-R3, but a different picture arose with F7-R7 primer set with the CST-Healthcare effluent having the highest *intl-*1 diversity.

#### 3.2.3 Laboratory Validation of selected *intI*1 primers to quantification *intI*1 mRNA transcript from environmental samples

As detection of *intl*1 DNA does not infer integrase activity, each of the validated primer sets were tested for their ability to quantify *intI*1 mRNA transcripts. For this, fresh river water samples were used as the quality of the RNA extracted wastewater nucleic acids maybe of poor quality due to long term storage(Cholet et al., 2019), although quality was not measured. For each primer set, the reverse transcriptase reaction was carried out with random hexamers (RH) and gene specific (GS) primers as previous work showed increased specificity with gene specific priming (Cholet et al., 2020). In addition, Q-PCR quantifications were carried out with (Figure 4C) and without the probes (i.e. SYBR Green) (Figure 4A, 4B, Table 4). All primer sets successfully quantified *intl*1 DNA and mRNA from river water, with *intl*1 gene abundances greater than *intl*1 transcripts (Figure 4).

**Figure 4:**
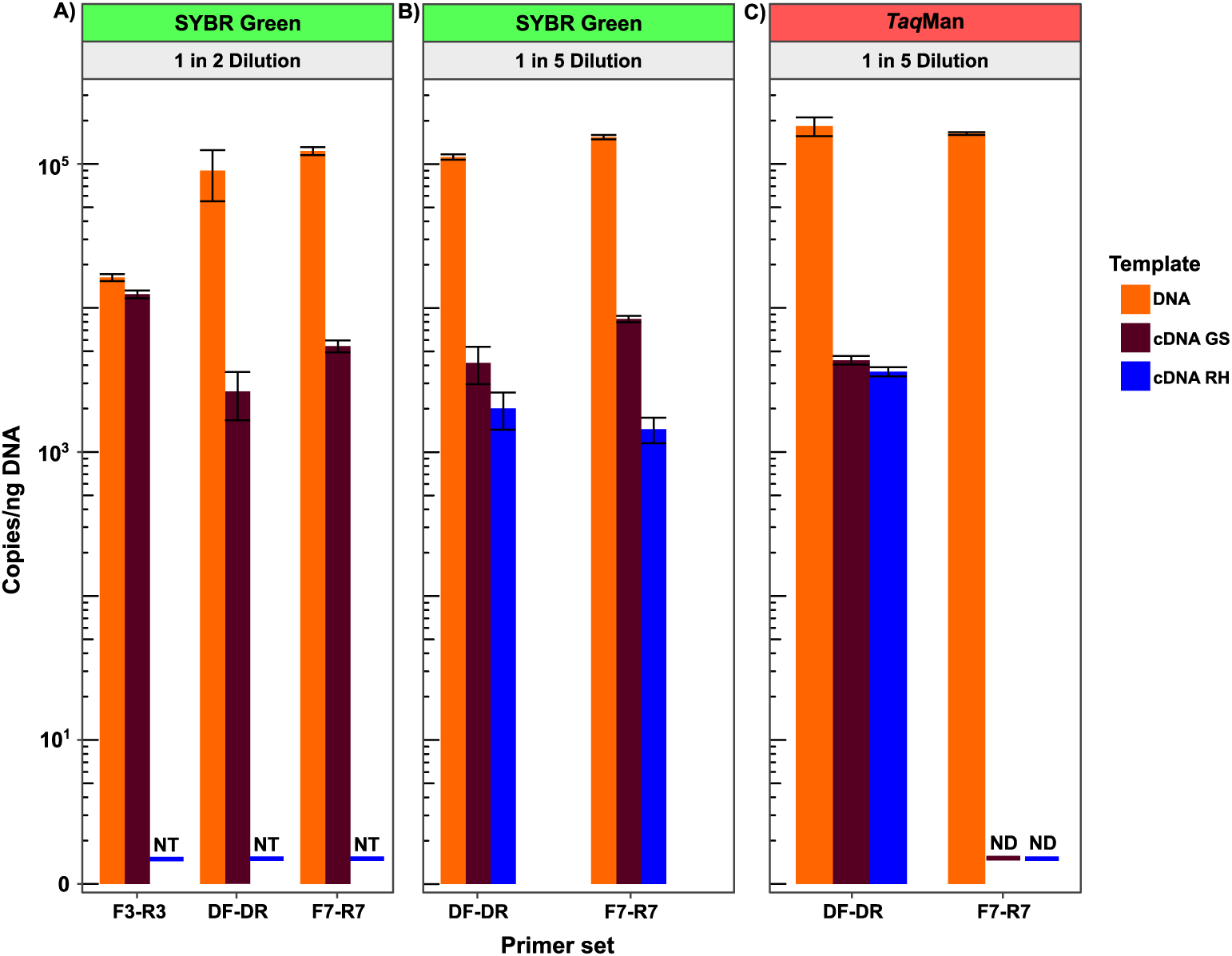
*intI*1 DNA and mRNA transcript quantified from river water sample by the three selected *intI*1 primer sets (DF-DR, F3-R3, F7-R7). Reverse transcriptase reaction for each primer set was performed with random hexamers (RH) and gene specific (GS) primers. Additionally, *Taq*Man assays were carried out with (C) and without the probes (i.e. SYBR Green) (A, B). NT denotes not-tested, ND denotes non-detected.

As previously shown (Cholet et al., 2020), gene specific priming was more efficient than random hexamer priming. F7-R7 primer set did not work as a *Taq*Man probe assay but worked in a SYBR green assay. It should noted that, whilst higher *intI*1 transcript copies per ng DNA were quantified by the F3-R3 SYBR Green assay (1 in 2 dilution), direct comparison to DF-DR and F7-R7 cannot be made as they were not done on the same sample. The aim here was simply to demonstrate that the primer sets were able to quantify *intI*1 mRNA transcripts. In summary, the primer sets tested are appropriate to quantify *intI*1 mRNA transcripts from environmental samples, using both gene specific and random hexamer priming, albeit that the *Taq*Man probe chemistry must be swapped to SYBR green chemistry if the F7-R7 primer set is to be used.

## 4.0 Discussion

Accurate quantitative data is key to inform evidence-based management strategies and policies to reduce the global AMR burden. Quantitative approaches, alongside unified methodologies to enable comparison among data sets is a powerful tool to enable this. The clinical class 1 integron (CL1-integron) integrase (*intI*1) has been proposed as a proxy for inferring potential AMR (Gillings et al., 2015). The first step to investigating the potential for this is to select appropriate primers, however our systematic literature review revealed over 65 *intl1* primer sets with little consensus on the best primer to use. Through *in silico* testing of the published primer sets, in addition to the design of an optimised primer set in this study, we selected three *intl1* primer sets for laboratory validation and further testing of their specificity on septic tanks from Thailand associated with healthcare and household usage to investigate their contribution in disseminating CL1-intgerons to the environment. This included a novel solar septic tank designed with internal heating ranging from 39 to 63.6°C in the disinfection chamber. From the 65 primers in the literature, three were selected-two published primer sets, F3-R3 (Rosewarne et al., 2010) and F7-R7 (Barraud et al., 2010) which have been extensively applied to survey CL1-integron abundance in a range of ecological settings including WWT (Stalder et al., 2014; Paiva et al., 2015) and agricultural settings (Johnson et al., 2016; McKinney et al., 2018) and a newly designed primer, DF-DR, modified from the F3-R3 primer set and an MGB probe added to increase specificity showed good coverage and specificity. All were successful PCR and RT-Q-PCR assay.

To confirm their specificity, MiSeq amplicon sequencing of the short amplicons was undertaken from the Thai septic tank samples. While diversity of the amplicons was low, likely reflecting the short amplicon length, the *intI*1 gene was ubiquitous in our samples supporting previous findings where it was dominant in polluted environments such as WWT (Gillings et al., 2015; Zheng et al., 2020). The ASVs generated from each primer sets were highly similar to *intI*1 and *intI*1-like sequences obtained from known and unknown bacteria, which were isolated from a range of clinical settings (e.g., human commensal) and environmental sources (e.g., wastewater activated sludge) (Figure 3). Interestingly, a few of the *intI*1-like sequence characterised were from known bacterial species isolated within clinical context including human faeces and bloodstream (Figure 3). This observation challenges the well-established knowledge that *intI*1 sequence recovered within clinical settings have identical/nearly identical protein (≥98% protein identity) (Roy et al., 2021) and/nucleotide sequence (99-100% nucleotide identity) (Gillings et al., 2008a) to each other. As such, this implies that *intI*1-like sequences can also be present within clinical settings and not just restricted to environmental settings as originally thought.

Our sequence results also highlight that the primer sets show that it was not possible to distinguish between *intl1* and the lesser conserved *intI*1 variants (*intI*1-like, <98% protein similarity) that has been shown to co-exist within these settings and similar environments (Gillings et al., 2008a, 2008b). These less conserved CL1-integron integrases (*intI*1-like) have been found, for example, on the chromosome of non-pathogenic *Betaproteobacteria* isolated from biofilms and soil and the entrained gene cassettes encoded currently unknown function rather than AMR (Gillings et al., 2008a). *intl*1-like may therefore not contribute to AMR but will contribute to *intl*1 Q-PCR signal. None of the primers, not even with the addition of a *Taq*Man or MGB probe, were able to distinguish between *intl*1 and *intl*1*-*like. As such, quantified *intI*1 gene abundance could potentially be overestimated. However, designing new primers over longer region but still suitable for QPCR, capable of distinguishing both variants can be a challenge. This is because *intI*1-like protein sequence identity between bacteria species can vary when compared to the reference *intI*1 *Pseudomonas aeruginosa* reference sequence (pVS1, AAA25857.1). For example, the *intI*1-like protein sequence from *Salmonella enterica subsp* (KMJ40944.1), a gamma-proteobacteria and *Comamonas thiooxydans* beta-proteobacteria (WP_012838479.1) shared 87.8% and 92.8% identity to the reference pVS1 *intI*1 protein sequence respectively. As such, the varying conserved region shared between the *intI*1, and *intI*1-like sequence variant makes it a challenge to design a primer that exclusively distinguish both variants.

The potential contributions of *intI*1-like abundance to overall abundance of *intI*1 gene quantified via Q-PCR suggests that *intI*1 abundance may not be an adequate or reliable proxy for inferring overall AMR abundance. Therefore, other potential proxies such as the *qacEΔ*1 (confer antiseptic resistance) or *VanA* (confer vancomycin resistance) (Abramova et al., 2022) should be investigate for reliable estimation of overall AMR abundance.in polluted environment.

This work has also shown the impact of using different primers on the interpretation of the findings and in turn our understanding of the risk of AMR. While across our septic tanks, the three best performing primers sets revealed the same overall trends (Figure 2), they did on occasion change the statistical difference between samples. For example, there were statistically higher *intl*1 gene abundances in the effluent than the sludge of the CST-healthcare unit when quantified with one (DF-DR) of the three primer sets but no difference when using the other two (F3-R3 and F7-R7) (Figure 2). Depending on the primer set used, our understanding of the role of wastewater in the dissemination of CL1-integron and entrained AMR gene to the wider environment differed, highlighting the need for primers standardisation if comparisons and environmental meaning are to be gained from the large body of literature and work currently being undertaken in this area. With this in mind, from the work carried out validating and comparing the primer sets, we arrived to three very good primer sets, albeit with the lack of specificity for *intl-1*. As the addition of the TaqMan and MGB probes did not offer increased specificity, we recommend F3-R3 primer set and SYBR green assay (Rosewarne et al., 2010). This primer set has previously been extensively used in the literature to survey CL1-integrons from a wide-ranging environment (Paiva et al., 2015) and here we have further demonstrated their suitability to quantify mRNA also. For this a gene specific RT-Q-PCR assay performed best as previously demonstrated (Cholet et al 2020).

### Ecological risk assessment of septic tanks in contributing to *intI*1 gene abundance to the environment

Comparing the abundance of *intI*1 genes (copies/ng DNA) among the different septic tanks, we showed that they were higher in the effluent compared to sludge, for all three reactors (CST-household, CST-healthcare, SST-household), irrespective of the *intI*1 primer set used, with the highest gene abundance quantified in the conventional healthcare (CST-healthcare) effluent (Figure 2). This finding was consistent with a previous study that reported higher *intI*1 relative gene abundance (normalised abundance to the *16S rRNA* copies) in hospital effluent compared to urban or municipal WWTP effluent (Stalder et al., 2014). Healthcare institutions are among the primary consumers of antimicrobials particularly antibiotics (Stalder et al., 2014). As such, stronger selective pressures are imposed within the bacteria communities, which in turn drives acquisition of resistance genes carried within key vector such as CL1-integron, to ensure their survival from the constant threat of antimicrobials within WWT system.

Of the three reactors, lower *intI*1 gene abundance per ng of DNA was quantified in the household solar septic tank (SST) samples (sludge and effluent) compared to the conventional tanks (CST-healthcare and CST-household) by two (DF-DR and F7-R7) of the primer sets whilst the third (F3-R3) only quantified lower *intI*1 gene copies in the SST-household effluent and not the sludge sample (Figure 2). Nonetheless, this implies that the increase temperature potentially plays a role in reducing CL1-integron from WWT and thus, the abundance entering the environment. This finding agrees with our proposed hypothesis of decreased *intl*1 gene abundance as a result of increased temperature driving enhanced wastewater treatment. Although the target internal temperature (50-60°C) within the solar tank was not consistently achieved, our finding is consistent with the recent study by Zhang and colleagues (Zhang et al., 2022), who investigated removal of CL1-integron and entrained AMR genes from anaerobic digestors operated at higher (thermophilic-55°C) and lower (mesophilic-35°C and 25°C) temperatures and reported statistically lower *intI*1 gene abundance and removal at higher temperature. In addition, statistically lower *16S rRNA* gene abundance was reported at the higher temperature, coupled with a lower relative abundance of AMR gene cassettes, albeit slightly higher ARG subtypes was detected with the higher temperature.

Although typified by poor treatment performance (Connelly et al., 2019), the conventional household tank was able to reduce *intI*1 gene abundance in the effluent from the influent by 31.21 to 42.33%, depending on the *intI*1 primer set used. This finding is consistent with previous study by Chen and Zhang (2013), although Chen and co-worker reported higher *intI*1 removal in the effluent from the influent (estimated around 1.9 to 2.3-log removal) for two of the three onsite domestic WWT associated with single family usage investigated than observed in this study. In the third onsite domestic WWT associated with single-family use enrichment of *intI*1 gene abundance was reported. However, the better removal from the two tanks in their study (Chen and Zhang, 2013) may be due to the additional secondary treatment incorporated to the tank, such as eco-filter, constructed wetland and multi-soil layering, prior to discharge to the environment which was not done in our study.

WWT sludge represents an additional source of CL1-integron and entrained AMR genes to the environment, particularly if improperly managed (i.e. improperly disposed of without further treatment), which further exacerbate global AMR burden (Koottatep et al., 2021). In the Global south region such as Thailand and Vietnam, only 10-20% of the faecal sludge generated are estimated to be adequately disposed of, whilst the vast majority are discharged directly to the environment (Koottatep et al., 2021). With the high *intI*1 abundance quantified in the sludge for the three reactors, coupled to the already high abundance in the effluent, we found that, on average, 1.22×10^5^ to 1.48×10^5^, 8.41×10^4^ to 1.1×10^5^, and 7.73×10^4^ to 9.4×10^4^ *intl*1 gene copies per ng DNA (depending on primer set), enters the environment via the CST-household, CST-healthcare and SST-household respectively. This is significant when taking into account the proportion of global population (2.7 billion people) estimated to be served by onsite decentralised WWT including septic tanks (Harada et al., 2016). Thus, highlighting septic tanks as an important source of CL1 to environment, and further supports the broader knowledge that WWT in general, are a major source of CL1-integrons and entrained resistance genes to the environment. For the CST-household tank with accessible influent sample, whilst the load of *int*I1 was decreased from the influent, the abundance *intI*1 quantified in the sludge and effluent by the different primer sets represent a significant source of *intl*1 to the environment and therefore, emphasise the need to optimise the conventional septic tank for AMR removal.

The increased abundance of CL1-intgerons entering the natural environment from WWT coupled to a slow decay rate (*intI*1 halve-life estimated ≥ 1 month in soil (Burch et al., 2014)), increases the risk of acquisition and dissemination into broader bacteria taxa especially clinically relevant human pathogenic bacteria including *Acinetobacter baumannii* (Nikibakhsh et al., 2021), *Proteus mirabilis* (Chen et al., 2017; Lu et al., 2022) and *Pseudomonas aeruginosa* (Liu et al., 2020; Khademi et al., 2021).

## 5.0 Conclusions

This present study has provided insight into the importance of primer choice especially in the context of validating the *intI*1 as a suitable proxy for AMR pollution, and the need for standardisation across studies to comprehensively understand the role in which wastewater play in disseminating CL1-intgerons and by extension AMR genes to the environment. Further work is needed to determine if the *intl*1 is indeed a suitable proxy for over overall AMR gene abundances.

Moreover, we showed septic tank decentralised wastewater, particularly the conventional healthcare tank (CST-healthcare), can be a significant source of CL1 integron to the environment via the effluent and sludge if the sludge is directly applied to the environment without undergoing additional treatments, such as wetlands, to reduce the *intI*1 gene load. Thus, supports growing evidence that WWT in general are a major source CL1-intgerons and associated resistance genes to the wider environment which further exacerbate the global burden from AMR.

## Acknowledgements

We gratefully acknowledge the support of the following funders: VO was supported through a University of Glasgow College of Science and Engineering Doctoral studentship and CS by a RAEng of Engineering-Scottish Water Research Chair (RSF1718943). The project was funded through EPSRC awards EP/VO30515/1 and EP/P029329/1.

## Notes

### Competing Interest Statement

The authors have declared no competing interest.

