## Supplementary materials for "*intI*1 primer selection for class 1 integron integrase gene and transcript quantification – validation and application for monitoring *intl*1 gene abundance within septic tanks in Thailand"

**List of supplementary Figure and Tables**

**Figure S.1:** Alignment of published and newly designed intI1 primers and probe sequence hit position to a *Pseudomonas aeruginosa* plasmid pVS1 nucleotide sequence (M73819.1).

**Figure S.2:** Description of synthetic gene *intI*1 fragment inserted into a circularised, double-stranded NGS verified plasmid vector used for constructing QPCR standard curve.

**Figure S.3:** Performance of intI1 primer sets against intI1 nucleotide sequences of SDB1 *(n= 104)* and SDB2 *(n=144)*, to evaluate primer coverage

**Figure S.4:** Performance of *intI*1 primer sets against intI1 nucleotide sequences of SDB3 (n= 502), to evaluate primer coverage.

**Figure S.5:** Performance of intI1 primer sets against *intI*1-like nucleotide sequences (*n= 15*) and non-*intI*1 nucleotide sequences (*n= 1540*), to evaluate primer specificity.

**Table S.1:** List of *intI*1 gene primer sets and probes reviewed in this study

**Table S.2:** Listed of *intI*1 primer sets excluded from further analysis in this study

**Table S.3:** Coverage and specificity of currently published and newly modified *intI*1 primer pairs

**Table S.4:** *intI*1 primer sets with no amplicon produced at 0 WS and the WS at which an amplicon was produced

**Table S.5:** Coverage of published and newly modified *intI*1 primer sets that incorporates a reporter probe

**Table S.6:** Two-way Anova test between primer sets for the same sample types

**Table S.7:** Summary statistics of the ASVs abundances per sample by MiSeq amplicon sequencing

**Supplementary Experimental Procedures**

**2.1 Primer evaluation**

Published *intI*1 primers were analysed as primer-pair (Table S.I), using Primer Prospector (Walters et al., 2011), to evaluate coverage and specificity against constructed integrase sub-databases (Figure 3). The analyze_primers.py function with the default settings on Primer Prospector was used to generate an alignment profile file for each primer against unaligned individual nucleotide sequence in each test sub-databases. For each primer alignment to a nucleotide sequence a weighted score (WS) was given.

The overall WS was calculated as:

Non-3’ mismatches * 0.4 per mismatch + 3’ mismatches * 1.0 per mismatch + Final 3’ base mismatch * 3.0 per mismatch + Non-3’ gaps * 1.0 per gap + 3’ gap * 3.0 per gap.

The first 5 bases of the primer and the target sequence were defined as the 3’end and thus, mismatches within these bases were termed 3’ mismatches. The remaining bases of the primer and the target sequence were defined as the non-3’ end. Therefore, mismatches within these non-3’ end bases were regarded as non-3’mismatches. Gaps in the alignment of the primer and the target sequence in the first 5 bases were termed 3’ gaps while gaps in the alignment for the remaining primer and template sequence were known as non-3’gaps. A higher WS was given for mismatches or gaps at the 3’end compared to mismatches or gaps at the non-3’ end. This is simply because mismatches or gaps at the 3’end region has a significant chance of affecting PCR amplification, whereas mismatches or gaps at the non-3’ end can be tolerated. As such, the lower the WS, the better the compatibility between the primer and target DNA sequence which suggest higher primer coverage potential. A 0 score indicates perfect alignment. Only sequences that had the correct primer-binding orientation (5’ to 3’ directionality) were analysed since DNA synthesis is from 5’ to 3’.

To evaluate primer coverage and specificity of each primer pair, the primer-binding orientation (i.e., reverse primer alignment was before the forward primer alignment position) against each unaligned nucleotide sequence was first verified from the primer hit position. Sequences with incorrect primer-binding orientation as a result of missing primer binding site were discarded from each sub-database. Following this, the number of amplicons estimated to be amplified by each primer pair was calculated using the sum of the WS of the forward and reverse primer for each primer pair. If the sum was ≤ a defined threshold (0 - perfect match to 10 – incompatible match), then the primer set was considered to amplify the target sequence within the test database. Furthermore, the mean overall WS for the forward and reverse primer of each primer set was calculated by taking the average of the WS for the total number of sequences with the correct binding orientation (Table S.III). Lastly, the R package “ggplot2” (Wickham, 2009) was then used to generate a WS plot of each primer set based on the defined WS threshold.

**2.2 Design of *Taq*Man-minor-groove binder (*Taq*Man-MGB) probe**

To design an *intI*1 MGB-TaqMan probe, *intI*1 sequences within SDB3 (*n=503*) were aligned using the MAFFT algorithm. Aligned sequences were imported into EMBOSS Cons website (Last accessed 04/08/2021 <https://www.ebi.ac.uk/Tools/msa/emboss_cons/>) to generate a consensus sequence. The consensus sequence was exported into Primer Express software (Version 3.0.1; Applied Biosystems)^TM^ with selection of *Taq*Man MGB Quantification option. MGB probe design parameter was set to a minimum length of 13 bp and a maximum of 15 bp. Probe sequence with minimal secondary structures, and closer to the position of the forward or reverse primer was selected. Modified *intI*1 primer set (DF-DR) and designed MGB *Taq*Man-probe sequence (Table S.I) were BLAST searched (BLASTN) to validate the sequence specificity before proceeding to the alignment of probe sequence to reference pSV1 *intI*1 gene sequence (Figure S.1) and subsequently, *in silico* validation across constructed integrase sub-databases as specified above (see section 2.1).

**2.3 QPCR standard curve construction**

Q-PCR standard curves were constructed by amplifying synthetic *intI*1 gene fragment containing the primer binding site for all selected primers inserted in a circularised, next-generation sequencing (NGS) verified, ampicillin-resistant vector from Integrated DNA Technologies (Figure S.2). Briefly, *Escherichia coli* (NC_011964.1) *intI*1 gene fragment containing the primer-binding site for the three selected primers, with additional 10 bases at the ends of the total primer site, was flanked with a T7-forward (5’-TAATACGACTCACTATAGGG-3’) and M13 reverse primers (5’-CAGGAAACAGCTATGAC-3’), resulting in a 660bp gene fragment (Figure S.2). The 660bp gene fragment (sequence can be found in below Figure S.2) was then inserted in a circularised, NGS verified, ampicillin-resistance vector (pUCIDT-AMP). pUCIDT-AMP vector DNA was resuspended in a 20µl IDTE (10mM Tris, 0.1mM EDTA) buffer at pH 7.8, with a final concentration of 200ng/μl.

PCR was carried out using T7 (5’-TAATACGACTCACTATAGGG-3’) and M13 (5’-CAGGAAACAGCTATGAC-3’) flanking primers, with the HotStartTaq PCR kit (Qiagen) in a 25μl volume, which consisted of 15.875μl nuclease-free water, 2.5μl 10x PCR Buffer, 0.125μl HotStartTaq, 0.5μl dNTPs (10μM), 0.5μl of each primer (10 μM each), and 5μl (10ng) template DNA (pUCIDT- vector). The reaction condition was as follow: 95°C‐15 min, (94°C‐30s, 57°C‐30s, 72°C‐60s) ×29 cycles and a final extension at 72°C for 10 mins. The resultant PCR product was cleaned, and size selected with the Agencourt AMPure XP beads.

**2.4 Sampling**

CST influent was sampled by disconnecting inflow to the CST septic tank via a sampling valve for 24-hours. Waste generated during the 24-hour period were collected in a sealed bucket, followed by homogenisation of the buckets’ content. Three 1 L homogenised samples were then collected in storage bottles and stored at -80^O^c if not in use immediately for downstream processing. The physiochemical and tanks operational parameters of all septic tank units were measured. Due to inaccessibility of influent samples, influent was only collected for one CST-household unit (CST-P3).

CST and SST effluent and sludge samples were collected prior to influent sampling to ensure samples were representative of the system under normal operating conditions. Sampling of effluent was done by flushing the toilet once to clear the outflow pipe of residual materials, followed by collection of effluent in a 10L bucket after a second flush. The effluent was homogenised by mixing, and three 1L sub-samples were collected in 1L bottles for later use. Conversely, sampling of sludge was done by homogenising the tank contents by mixing using a submersible pump. Subsequently, 2L of homogenised sample was collected into a plastic beaker through tubing (2cm internal diameter) inserted to the tank and connected to an external vacuum pump (Sacco, Model SC-1A). Contents of the beaker were thoroughly mixed by stirring and then four sub-samples taken in 50mL centrifuge tubes before storing on ice (approximately 2 hours) and transported to the laboratory for downstream processing.

**2.5 MiSeq Amplicon sequencing**

The specificity of the selected *intI*1 primer sets (DF-DR, F3-R3, F7-R7) used to quantify *intI*1 gene from 31 septic tank wastewater, were confirmed by Illumina MiSeq amplicon sequencing. A two-step PCR was performed to barcode samples as detailed previously(Bourlat et al., 2016; Cholet et al., 2019). To do so, a two-step PCR was performed using a similar method detailed previously (Bourlat et al., 2016; Cholet et al., 2019). The first PCR step amplified the target region using respective *intI*1 primers (primer sequences outlined in **Table III**), attached with Illumina adaptors at the 5’ end: 5′‐TCG TCG GCA GCG TCA GAT GTG TAT AAG AGA CAG- 3’ (forward adaptor); 5′‐GTC TCG TGG GCT CGG AGA TGT GTA TAA GAG ACA G- 3’ (reverse adaptor). For each primer set, PCR amplification was carried out in a 25μl volume reaction using 5μl (5ng) template DNA and the HotStartTaq PCR kit (Qiagen). Each 25μl volume reaction consisted of: 15.75μl (DF-R7 and F3-R3)/ 14.75μl (F7-R7) nuclease-free water, 0.5μl (DF-R7 and F3-R3)/ 1μl (F7-R7) of each primer (10 μM each), 0.5μl dNTPs (10 μM each), 0.25μl HotStartTaq and 2.5μl of 10x PCR Buffer.

A no template control and positive control (plasmid vector containing target sequence) were included for each primer set. The PCR thermocycling conditions are specified in **Table III**. Gel electrophoresis was performed on generated amplicons to confirm the expected amplicon size and quality. For each primer set, duplicate, or triplicate PCRs were carried out on the samples (depending on the intensity of the band seen on gel following gel image analysis), using the same conditions, and then pooled together for further processing. PCR amplicons were cleaned, and size selected using a 1.5X volume ratio Agencourt AMPure XP beads (Beckman Coulter, Brea, CA, USA) according to the manufacturer's recommendation and eluted in 30μl of nuclease-free water.

The second PCR step (index PCR) was performed to incorporate Illumina dual index (i5 and i7) using the Nextera XT Index Kit in a 25μl volume reaction which consisted of: 6.75μl nuclease-free water, 2.5μl of 10x PCR Buffer (Qiagen), 0.25μl HotStartTaq, 0.5μl dNTPs (10 μM each), 5μl of each index primer (10 μM each), and 5μl template. The cycle conditions are detailed in **Table III**. Amplicons generated were purified using a 1.5X volume ratio Agencourt AMPure XP beads and eluted in 25μl nuclease-free water. Following this, three samples were chosen at random from each primer set and ran on the Bioanalyser following the DNA 1000 Assay protocol (Agilent Technologies, UK) to determine the average length of the amplicons generated by each primer set, and to confirm the absence of unspecific products. The DNA concentration of each amplicon, from each primer set, was determined fluorometrically (Qubit) and molarity was calculated using the following equation:

(Concentration in ng/μl) × 106 = (660 g/mol × average library size)

For each primer set, prepared libraries were pooled at an equimolar amount into individual tubes. Subsequently, the three libraries were pooled at an equimolar amount to make the final pooled library. Lastly, the final pooled library was measured fluorometrically (Qubit) before sending to GENEWIZ (GENEWIZ Sequencing, UK) for MiSeq amplicon sequencing on the Illumina platform (2 × 250 bp paired-end).

**2.6 DNA/RNA Co-extraction**

All glassware was baked at 180°C overnight to inactivate RNases. Lids of glassware and stirrers were soaked overnight in RNase Zap (Ambion) and disposable plasticwares used, including tubes, were RNase free. All solutions used for nucleic acid extraction were prepared using diethylpyrocarbonate (DEPC) treated Milli-Q water. Total nucleic acid using the phenol-chloroform method described by Griffiths et al., (2000), with a minor modification to the bead-beating time as outlined by Lim et al., (2016). Briefly, glass microfibre filters were split in halves using sterile forceps, and each half placed in a matrix E-bead-beating tubes (MP Biomedical) and immediately transferred on ice. 0.5ml 5% CTAB (hexadecyltrimethylammonium bromide)/phosphate buffer (120mM, pH8; consisting of 2.58g K2HPO4.3H2O; 0.10g KH2PO4; 5.0g CTAB; 2.05g NaCl; 100ml DEPC water) and 0.5ml Phenol:Chlorophorm:Isoamyl alcohol (25:24:1; v:v:v; pH8) were added to each bead-beating tube. Samples lysed on the FastPrep system (MP Biomedical) at 6.0m s^-1^ for 45s, and then centrifuged at 12,000g for 20 mins at 4°C. The top aqueous layer was transferred to a sterile 2ml tube and mixed with 0.5 ml chloroform:isoamyl alcohol (24:1 v:v) followed by a centrifugation at 16,000 g for 5 mins at 5^O^C. The top aqueous layer was transferred to a new sterile 2 ml tube and total nucleic acids were precipitated by adding 2 volumes of 30% polyetlyleneglycol 6000 (PEG6000)/ NaCl (1.6M) solution to the tube. The resulting mixture was incubated on ice for 2 hours and then centrifuged at 16,000g for 30 mins at 4°C. The pellet was carefully recovered, by discarding the supernatant, and then washed with 1ml ice-cold 70% ethanol, followed by centrifugation at 16,000g for 30 mins at 4^°^C. The ethanol wash was carefully discarded, and the tube (containing pellet) was spun briefly for 5 sec at 4°C to remove residual ethanol. Recovered total nucleic acids pellet was air dried and subsequently re-suspended in 50μl DEPC treated water. Concentration of DNA in samples were determined fluorometrically (Qubit) and gel electrophoresis was performed to confirm success for DNA and RNA co-extraction. Co-extracts (DNA and RNA) were stored at -80˚c if not used immediately.

**2.7 RNA preparation and cDNA synthesis**

RNA was prepared from the raw DNA/RNA co-extract by DNase treating with Turbo DNase Kit (Ambion) in accordance with the manufacturer’s recommendation, with modification to the incubation time and volume of DNase added. 1μl DNase volume was added to the samples and incubated at 37^°^C for 1 hour, followed by further addition of 1μl DNase volume to the sample and a re-incubation at 37^°^C for another hour. Subsequently, the success of DNase treatment was confirmed by no PCR amplification of the V4 - V5 region bacterial 16S rRNA gene using the 515F (5’-GTGYCAGCMGCCGCGGTAA-3’) and 926R (5’-CCGYCAATTYMTTTRAGTTT-3’) primers (Suzuki et al., 2000). The PCR amplification was carried out in a 25µl volume reaction with the HotStartTaq PCR kit (Qiagen) containing 18.8μl nuclease-free water, 2.5μl 10x PCR Buffer, 0.2μl HotStartTaq, 0.5μl dNTPs (10μM), and 0.5μl of each primer (10μM each) and 2µl neat template (DNase treated RNA). The reaction condition was as follow: 95^°^C ‐15 min, (94^°^C ‐45s, 50^°^C ‐30s, 72^°^C ‐40s) × 35 cycles and a final extension at 72^°^C for 10 mins. DNA-free total RNA concentration in the samples were quantified fluorometrically (Qubit) and by Bioanalyser following the RNA 6000 Nano Assay protocol (Agilent Technologies, UK). The RNA integrity number (RIN) was also determined by the Bioanalyser based on the 23S/16S rRNA ratio.

DNA-Free RNA was immediately reverse transcribed using the superscript IV reverse transcription kit (Invitrogen). Both Random hexamer (RH) and gene specific (GS) reverse transcription were performed for the DF-DR and F7_R7 primer sets and only GS reserve transcription for the F3-R3 primer set. The initial reverse transcription reaction mix which consisted of 8μl water, 1μl primer (10μM gene specific/ 50μM random hexamer), 1μl dNTP's (10μM each) and 3μl RNA template was incubated at 65^°^C for 5 min and immediately transferred to ice for 1 min. A second reaction mix which contained 4μl 5X first-strand buffer, 1μl 0.1 mM dithiothreitol (DTT), 1μl RNAse inhibitor (40 unit/μl) and 1μl SuperScript IV (200 unit/μl) was subsequently added and then incubated at 55^°^C for 10 min and 80^°^C for 10 min for gene specific priming/ 23^°^C for 10 min, 55^°^C for 10 min and 80^°^C for 10 min for Random hexamer priming (Cholet et al., 2019).

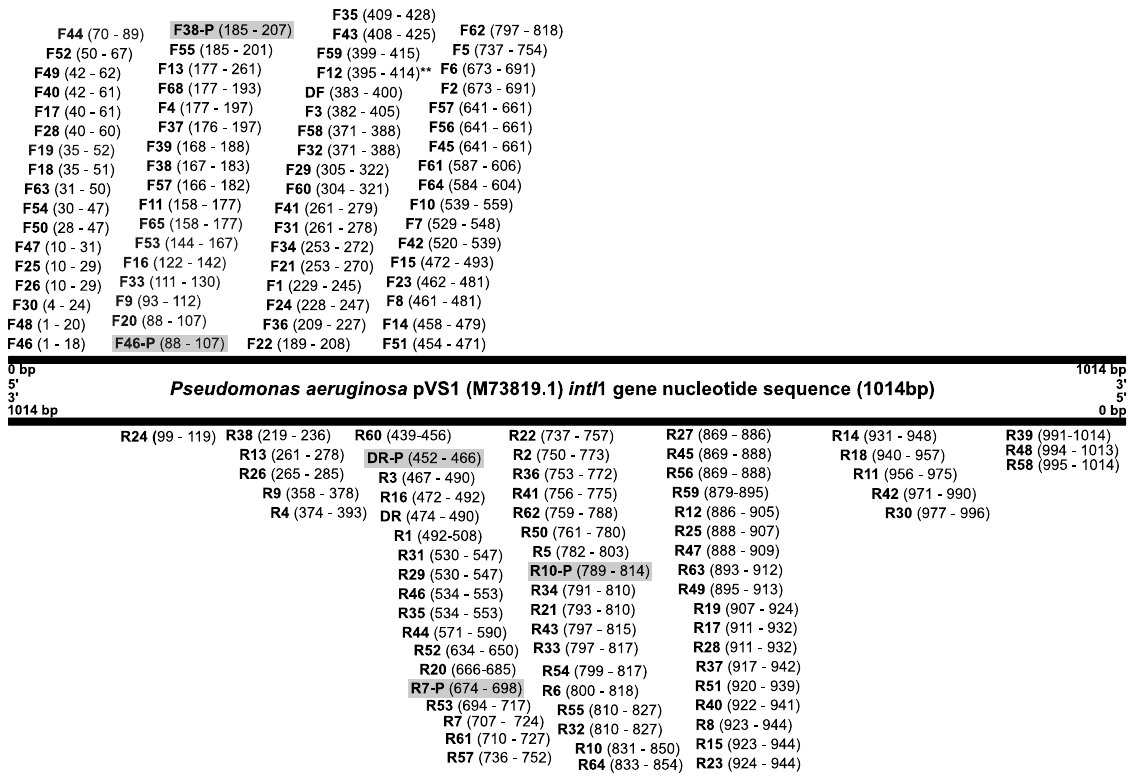

**Figure S.1:** Alignment of published and newly designed intI1 primers and probe sequence hit position to a *Pseudomonas aeruginosa* plasmid pVS1 nucleotide sequence (M73819.1). F refers to forward primer. R refers to reverse primer. The number after F or R (i.e. F1 or R5) refers to the assigned primer ID- See Table S.1 for more detail. The number in parenthesis () denotes the position of the primer sequence on the reference Escherichia coli intI1 gene. *** denotes primer hit position based on Primer Prospector alignment to CP003684.1 intI1 nucleotide sequence. Highlighted in grey are the probe binding position for primer-probe primer sets.

**660 bp insert fragment**

**T7 F. Primer**

**17 bp**

***intI1 gene insert (623 bp)***

**122	724**

***Escherichia coli intI1 gene (NC_011964.1)***

**Primer binding site (603 bp)**

**10 bp**

**5’**

**M13 R. Primer**

**20 bp**

**3’**

**10 bp**

**Figure S.2** Description of synthetic gene *intI*1 fragment inserted into a circularised, double-stranded NGS verified plasmid vector used for constructing QPCR standard curve. A 603bp region on *Escherichia coli* *intI1* sequence (NC_011964.1) containing the binding site for the selected *intI*1 primers for laboratory validation was flanked with extra 10 bases (total *E.coli* *intI1* gene sequence= 623 bases).The 623 bp *intI1* gene fragments was subsequently flanked with the T7-Foward and M13-Reverse primer. Total insert fragment= 660bp. The 660bp gene fragment was then inserted into an NGS verified circular, ampicillin resistant plasmid vector. F denotes forward primer and R reverse primer. The number following the forward primer indicates the hit start position of the first base of the forward primer while the number following the reverse primer indicates the hit position of the last base of the reverse primer. See Figure S.1 for detailed information of the binding position of the primers. The 660bp insert fragment sequence was:

>NC_011964.1 *Escherichia coli* plasmid insert fragment

TAATACGACTCACTATAGGGGTCCACTGGGTTCGTGCCTTCATCCGTTTCCACGGTGTGCGTCACCCGGCAACCTTGGGCAGCAGCGAAGTCGAGGCATTTCTGTCCTGGCTGGCGAACGAGCGCAAGGTTTCGGTCTCCACGCATCGTCAGGCATTGGCGGCCTTGCTGTTCTTCTACGGCAAGGTGCTGTGCACGGATCTGCCCTGGCTTCAGGAGATCGGAAGACCTCGGCCGTCGCGGCGCTTGCCGGTGGTGCTGACCCCGGATGAAGTGGTTCGCATCCTCGGTTTTCTGGAAGGCGAGCATCGTTTGTTCGCCCAGCTTCTGTATGGAACGGGCATGCGGATCAGTGAGGGTTTGCAACTGCGGGTCAAGGATCTGGATTTCGATCACGGCACGATCATCGTGCGGGAGGGCAAGGGCTCCAAGGATCGGGCCTTGATGTTACCCGAGAGCTTGGCACCCAGCCTGCGCGAGCAGCTGTCGCGTGCACGGGCATGGTGGCTGAAGGACCAGGCCGAGGGCCGCAGCGGCGTTGCGCTTCCCGACGCCCTTGAGCGGAAGTATCCGCGCGCCGGGCATTCCTGGCCGTGGTTCTGGGTTTTTGCGCAGCACACGCATTCGACCGATCCACGGAGCGGGTCATAGCTGTTTCCTG

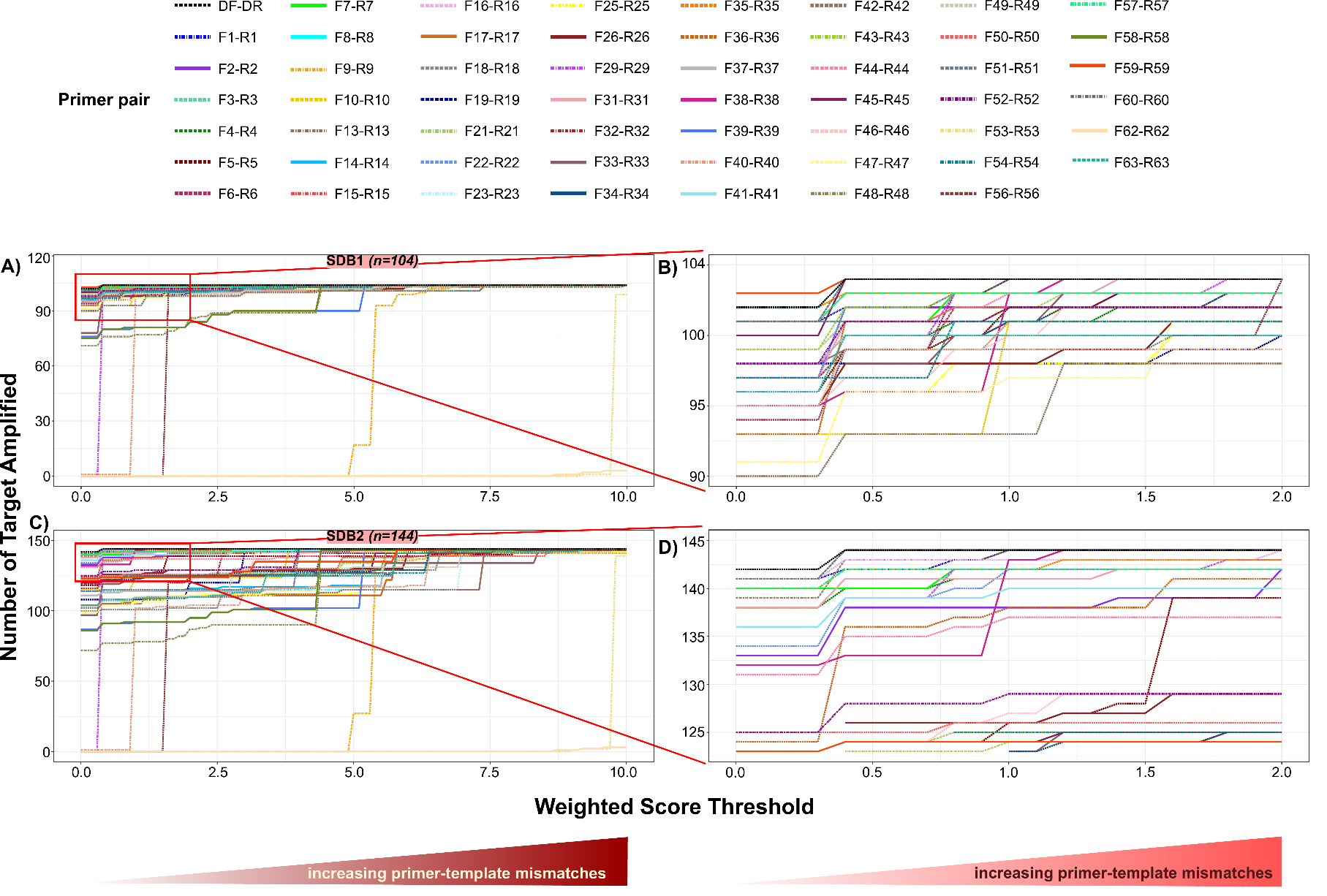

**Figure S.3:** Performance of intI1 primer sets against intI1 nucleotide sequences of SDB1 (n= 104) and SDB2 (n=144), to evaluate primer coverage. Primers were evaluated as pairs, for their ability to generate amplicon based on defined weighted score (WS) threshold that varied from 0 (strict) to 10 (less stringent). A WS of 0 indicates a perfect match (0 mismatch) between primer and template sequence. A WS >0 indicates mismatches between primer and template sequence. The top performing primers were defined as those primer set that were able to generate the highest number of amplicons at 0 WS in the test sub-database. A) SDB1 and C) SBD2 WS plot for all evaluated primer sets based on WS threshold that varied from 0 to 10. Red rectangular box indicate zoom in area of WS plot for SDB1 B) and SBB2 D). Each line colour and line type represent different set of primer.

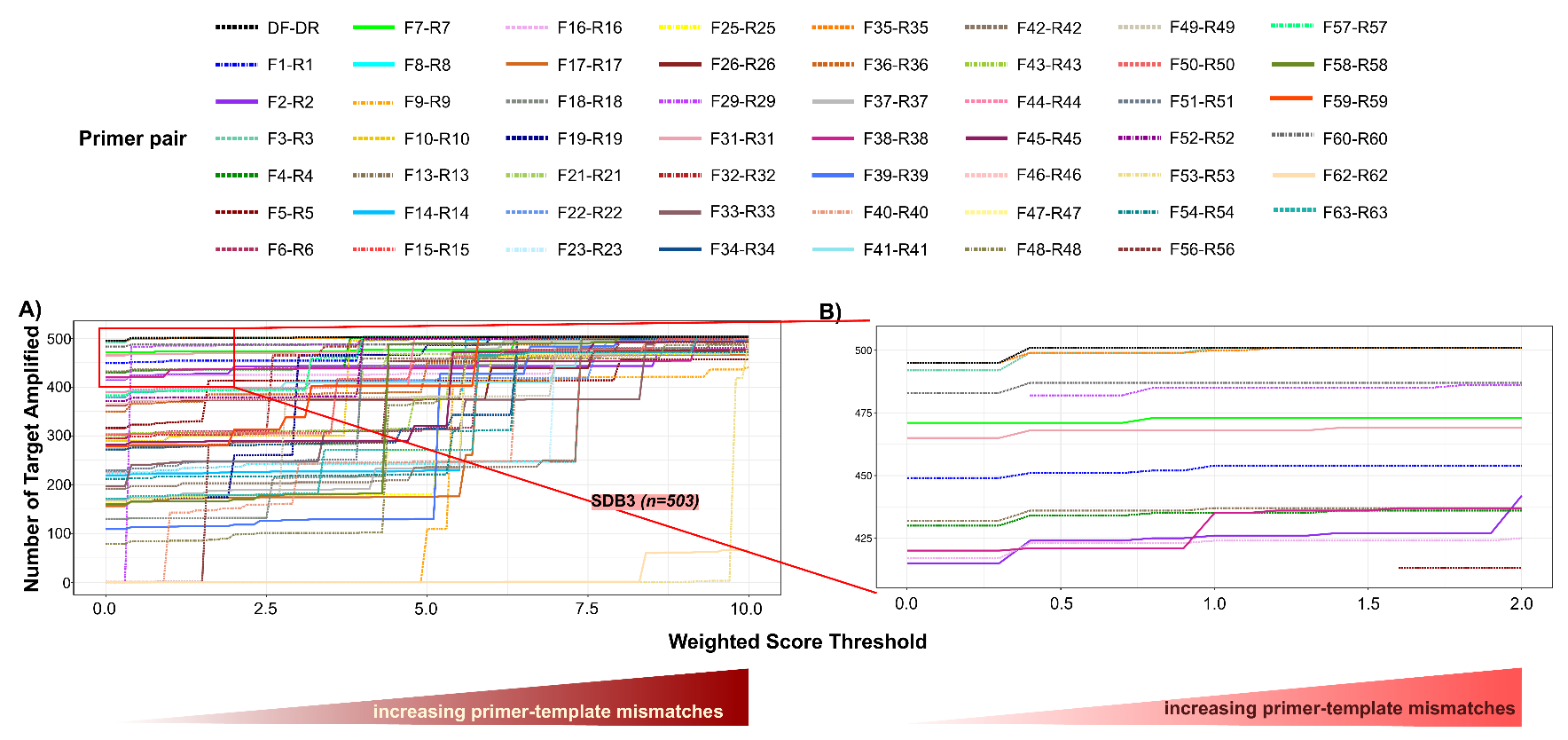

**Figure S.4:** Performance of intI1 primer sets against intI1 nucleotide sequences of SDB3 (n= 503), to evaluate primer coverage. Primers were evaluated as pairs, for their ability to generate amplicon based on defined weighted score (WS) threshold that varied from 0 (strict) to 10 (less stringent). A WS of 0 indicates a perfect match (0 mismatch) between primer and template sequence. A WS >0 indicates mismatches between primer and template sequence. The top performing primers were defined as those primer set that were able to generate the highest number of amplicons at 0 WS in the test sub-database. A) WS plot for all evaluated primer sets based on WS threshold that varied from 0 to 10. Red rectangular box indicate zoom in area. B) WS plot representing zoomed area of plot A. Each line colour and line type represent different set of primer.

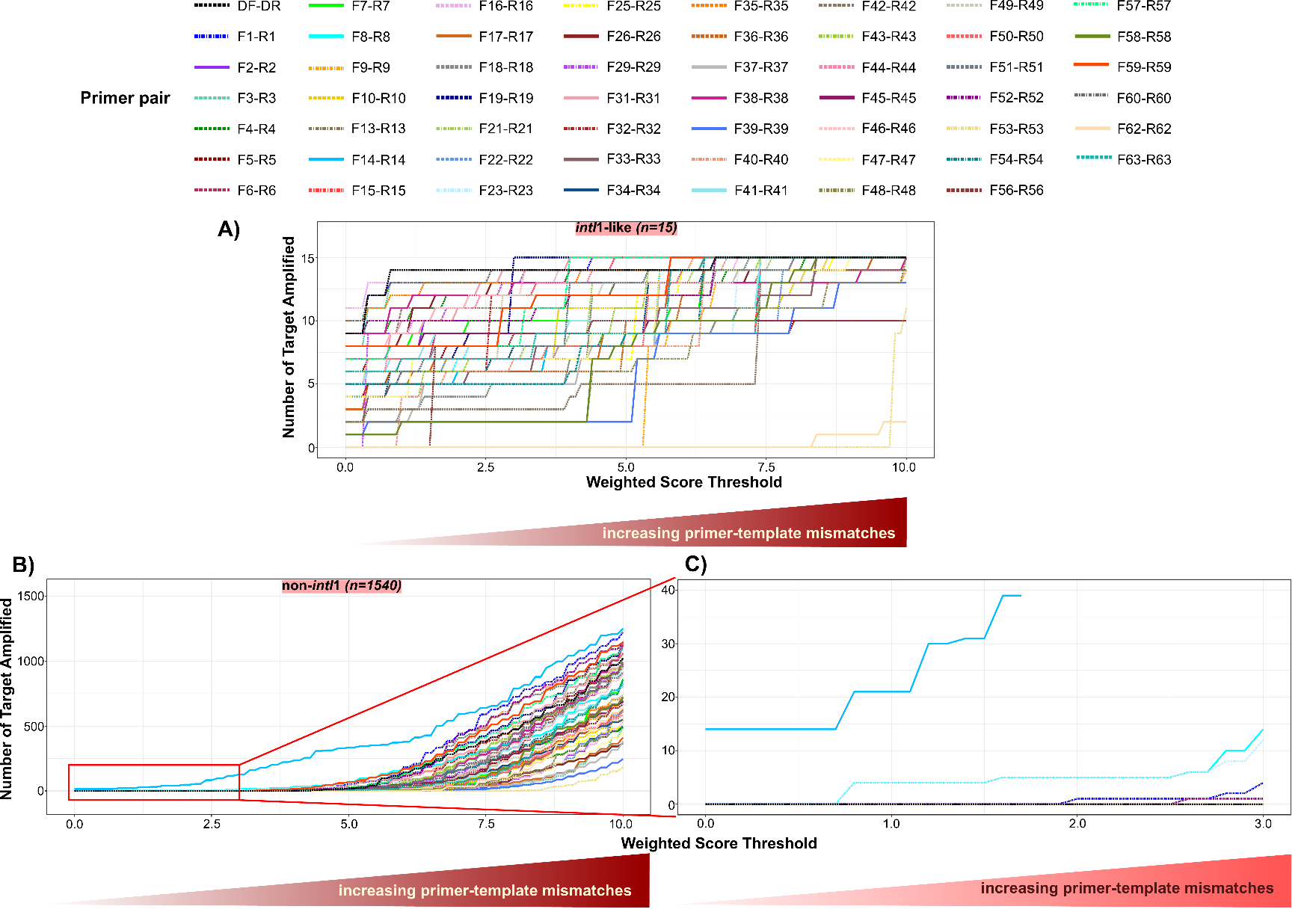

**Figure S.5:** Performance of *intI*1 primer sets against *intI*1-like nucleotide sequences (*n= 15*) and non-*intI*1 nucleotide sequences (*n= 1540*), to evaluate primer specificity. Primers were evaluated as pairs, for their ability to generate amplicon based on defined weighted score (WS) threshold that varied from 0 (strict) to 10 (less stringent). A WS of 0 indicates a perfect match (0 mismatch) between primer and template sequence. A WS >0 indicates mismatches between primer and template sequence. A) *intI*1-like and B) non-*intI*1-like WS plot for all evaluated primer sets based on WS threshold that varied from 0 to 10. Red rectangular box indicate zoom in area of WS plot for the non-*intI*1 sub-database C). . Each line colour and line type represent different set of primer.

**Table S.1:** List of intI1 gene primer sets and probes reviewed in this study

| **Primer source ID** | **Assigned ID -Position*** | **Sequence (5’-3’ Direction)** | **Primer length (bp)** | **Amplicon size (bp)** | **QPCR Study (QPCR Chemistry)** | **PCR Cycle Conditions** | **Environment** | **Reference** |
| --- | --- | --- | --- | --- | --- | --- | --- | --- |
| *intI1*-DF  *intI1*-DR  *intI1*-MGB | DF- 383  DR- 474  *DF_*P- 452 | TTCTGGAAGGCGAGCATC  TGCCGTGATCGAAATCC  Fam-TGACCCGCAGTTGCA-MGB | 18  17  15 | 108 | Yes  (MGB-TaqMan) | 95^O^ – 10min; (95^O^–30sec; 60 ^O^–60sec) x45 | Wastewater samples (Influent, sludge, and effluent) | **This study, modified from Rosewarne et al., 2010** |
| *intI1*/R  *intI1*/F | F1- 229  R1- 492 | TCCACGCATCGTCAGGC  CCTCCCGCACGATGATC | 17  17 | 280 | Yes (SYBR green I) | (94^O^–30sec; 55 ^O^–60^O^sec; 72^0^-15sec) x30 | Clinical isolates (Avian) | (Bass et al., 1999) |
| 345 R  245 F | F2- 673  R2- 750 | CATTCCTGGCCGTGGTTCT  TGAAAGGTCTGGTCATACATGTGA | 19  24 | 101 | Yes (SYBR green) | 95^O^C – 10min; (95^O^ – 15sec; 60 ^O^ – 60sec) x30 | Clinical isolates (Healthy adults) | (Skurnik et al., 2005) |
| qINT-4  qINT-3 | F3- 382  R3- 467 | TTTCTGGAAGGCGAGCATCGTTTG  TGCCGTGATCGAAATCCAGATCCT | 24  24 | 109 | - | 95^O^–10min; (95^O^-15sec ; 60^O^-15sec) x40 | Environmental samples (Sediments) | (Rosewarne et al., 2010) |
| *intI1*-a-Fw  *intI1*-a-RV | F4- 177  R4- 374 | CGAAGTCGAGGCATTTCTGTC  GCCTTCCAGAAAACCGAGGA | 21  20 | 217 | Yes (SYBR Green) | 95^O^–7min; (95^O^-10sec; 60^O^–30sec) x40; Melt curve analysis | Environmental samples (Sediment from fish farms) | (Muziasari et al., 2014) |
| Int1F2  Int1R2 | F5- 737  R5- 782 | TCGTGCGTCGCCATCACA  GCTTGTTCTACGGCACGTTTGA | 18  22 | 67 | Yes  (SYBR Green) | (50^O^-120sec; 95^O^-10min) x1; (95^O^-20sec ; 60^O^-60sec) x40; Dissociation curve step | Environmental samples (Industrial waste, sewage sludge and pig slurry) | (Gaze et al., 2011) |
| *intI1*-a-RV  *intI1*-a-Fw | F6- 673  R6- 800 | CATTCCTGGCCGTGGTTCT  GGCTTCGTGATGCCTGCTT | 19  19 | 146 | Yes (SYBR Green) | 95^O^–15min; (95^O^–15sec; 55^O^–30sec; 72^O^– 30sec) x45; melt curve: 65°to95° (0.5° increment/ 5sec) | Environmental samples (Surface water and sediment) | (Luo et al., 2010) |
| *intI1*-LC1  *intI1*-LC5  *intI1*-probe | F7- 529  R7- 707  F7-P- 674 | GCCTTGATGTTACCCGAGAG  GATCGGTCGAATGCGTGT  6 Fam- ATTCCTGGCCGTGGTTCTGGGTTTT-BHQ1 | 20  18  25 | 196 | Yes  (TaqMan probe) | 95^O^–10 min; (95^O^–30sec; 60^O^–60sec) x45 | Clinical /Laboratory strains | (Barraud et al., 2010) |
| *intI1*.F  *intI1*.R | F8- 461  R8- 923 | GGGTCAAGGATCTGGATTTCG  ACATGCGTGTAAATCATCGTCG | 21  22 | 483 | Yes (SYBR Green) | 94^O^–5min; (94^O^–30sec; 60^O^–30sec; 72^O^–60sec) x30 | Laboratory reference strains | (Mazel et al., 2000) |
| q*IntI1*F  q*IntI1*R | F9- 93  R9- 358 | ACCAACCGAACAGGCTTATG  GAGGATGCGAACCACTTCCAT | 20  21 | 286 | Yes (SYBR Green) | 95^O^–15min; (95^O^–30sec; 62^O^–30sec; 72^O^–30sec) x40; (72^O^–10min) x1; Melt curve analysis | Environmental samples (Riverine estuarine, and Freshwater) | (Wright et al., 2008) |
| *intI1*R476  *intI1*F165 | F10- 539  R10- 831  F10-P- 674 | TACCCGAGAGCTTGGCACCCA  CGAACGAGTGGCGGAGGGTG  Texas615-TCGTGATGCCTGCTTGTTCTACGGCA | 21  20  26 | 312 | Yes (*Taq*Man Probe) | 95^O^–5min; (95^O^–30sec; 62^O^–30sec; 72^O^–45sec; plate read) x40 | Environmental (Hospital, communal and urban WWTPs) | (Gillings et al., 2015; Paulus et al., 2019) |
| *IntI1*-R  *IntI1*-F | F11- 158  R11- 956 | CTTCAGCCTTTTCCAGCAAC  GAAACCTGCTCCAGCACTTC | 20  20 | 308  818*** | No | 94^O^–3min; (95^O^–30sec; 58.3^O^–30sec; 72^O^–40sec) x35; 72^O^–7min | Clinical isolates (Urine, blood, and wound) | (Najafgholizadeh Pirzaman and Mojtahedi, 2019) |
| intI-1F  intI-1R | F12- 395**  R12- 886** | TCATGGCTTGTTATGACTGT  GTAGGGCTTATTATGCACGC | 20  20 | 600  512*** | No | 94^O^–10min; (94^O^–40sec; 57^O^–50sec; 72^O^–55sec) x30-40; 72^O^–10min | Clinical isolates (Blood, urine, and wound) | (Mobaraki et al., 2018) |
| ZANi1F  ZANi1 | F13- 177  R13- 261 | CGAAGTCGAGGCATTTC  ACCTTGCCGTAGAAGAAC | 17  18 | 102 | Yes (SYBR Green) | 95^O^–3min; (95^O^–20sec; 57.8^O^–45sec; 72^O^–30sec) x40; 72^O^–10min | Environmental (Activated sludge, Pig Faeces) | (Yang et al., 2021) |
| hep35  hep36 | F14- 458  R14- 931 | TGCGGGTYAARGATBTKGATTT  CARCACATGCGTRTARAT | 22  18 | 491 | No | (94^O^–30sec; 55^O^–30sec; 72^O^–45sec) x30 | Clinical isolates (Urinary tract) | (White et al., 2000) |
| HS463a  HS464 | F15- 472  R15- 923 | CTGGATTTCGATCACGGCACG  ACATGCGTGTAAATCATCGTCG | 21  22 | 473 | Yes (SYBR Green I) | 94^O^–3min; (94^O^–30sec ; 60^O^–30sec; 72^O^–60sec) x35; Fluorescence acquisition step at 80^O^ | Environmental samples (Sediment) | (Hardwick et al., 2008) |
| HS916  HS915 | F16- 122  R16- 472 | TTCGTGCCTTCATCCGTTTCC  CGTGCCGTGATCGAAATCCAG | 21  21 | 371 | No | 94^O^–3min; (94^O^–30sec; 60/65^O^–30sec; 72^O^–90sec) x35; 72^O^–5min | Clinical isolates (UTIs from outpatients). | (Márquez et al., 2008) |
| intB  intA | F17- 40  R17- 911 | GTCAAGGTTCTGGACCAGTTGC  ATCATCGTCGTAGAGACGTCGG | 22  22 | 892 | No | (96^O^–5min; 55^O^–1min; 70^O^–3min) x1; (96^O^–15sec; 55^O^–30sec; 70^O^–3min) x24; 70^O^–5min | Environmental isolates (Estuarine environment) | (Rosser and Young, 1999) |
| *intI1*U/F  *intI1*D/R | F18- 35  R18- 940 | GTTCGGTCAAGGTTCTG  GCCAACTTTCAGCACATG | 17  18 | 923 | No | 94^O^–5min; (94^O^–30sec; 50^O^–30sec ; 72^O^–90sec) x30; 72^O^–7min | Clinical isolates (Healthy human patients) | (Zhang et al., 2004) |
| *intI1*F  *intI1*R | F19- 35  R19- 907 | GTTCGGTCAAGGTTCTGG  CGTAGAGACGTCGGAATG | 18  18 | 890 | No | 95^O^–2min; (95^O^–20sec; 54^O^–30sec; 70^O^–30sec) x35 | Environmental isolate (Soil, wastewater) | (Xu et al., 2007) |
| *intI1*F  Intf2 | F20- 88  R20- 666 | AGCTTACGAACCGAACAGGC  TCCGCCAGGATTGACTTGCG | 20  20 | 950 | No | 94^O^–5min; (92^O^–30sec; 55^O^–30sec; 72^O^–1min) x35; 72^O^–10min | Environmental samples (Sediments) | (Borruso et al., 2016) |
| intI-F  intI-B | **F21**- 253  **R21**- 793 | GCCTTGCTGTTCTTCTAC  GATGCCTGCTTGTTCTAC | 18  18 | 558 | No | 94^O^–5min; (94^O^–30sec; 55^O^–30sec; 72^O^–2.5min) x35; 72^O^–5min | Clinical/Reference isolates | (Guerra et al., 2001) |
| *intI1*R  *intI1*L | **F22**- 189  **R22**- 737 | ATTTCTGTCCTGGCTGGCGA  ACATGTGATGGCGACGCACGA | 20  21 | 568 | No | - | Clinical isolates | (Ploy et al., 2000) |
| *IntI1*-F  *IntI1*-R | **F23**- 462  R23- 924 | GGTCAAGGATCTGGATTTCG  ACATGCGTGTAAATCATCGTC | 20  21 | 483 | No | 94^O^–12min; (94^O^–30sec; 62^O^–30sec ; 72^O^–1min) x30; 72^O^–8min | Clinical isolates | (Machado et al., 2005) |
| *IntI1*F  *IntI1*R | **F24**- **228**  **R24**- 99 | CCCGAGGCATAGACTGTA  CAGTGGACATAAGCCTGTTC | 18  20 | 160 | No | (94^O^–30sec; 55^O^–30sec; 72^O^–30sec) x35 | Clinical isolates (including respiratory tracts) | (Koeleman et al., 2001) |
| Int-F  Int-B | **F25**- 10  **R25-** 888 | GCCACTGCGCCGTTACCACC  GGCCGAGCAGATCCTGCACG | 20  20 | 898 | No | 94^O^– 5min; (94^O^–15sec; 69^O^–30sec; 72^O^–60sec) x30; 72^O^–7min | Clinical isolates (UTIs strains) | (Kerrn et al., 2002) |
| Int-F  Int1-285B | **F26**- 10  **R26**- 265 | GCCACTGCGCCGTTACCACC  GCACAGCACCTTGCCGTAGAA | 20  21 | 276 | No | - | Clinical/ Environmental isolates (human, animal faeces and urine) | (Ho et al., 2012) |
| Inti1F  Inti1R | **F27**-182**  **R27**-869 | CGAGGCATAGACTGTAC  TTCGAATGTCGTAACCGC | 18  17 | 925 | No | - | Clinical isolates (Stool, blood, and urine) | (Orman et al., 2002) |
| Int1 R  Int1 F | F28- 40  R28- 911 | GTCAAGGTTCTGGACCAGTTG  ATCATCGTCGTAGAGACGTCGG | 22  21 | 550  892*** |  | 94^O^–5min; (94^O^–1min; 50^O^–1min; 72^O^–1min) x30; 72^O^–5min | Clinical isolates (Urine) | (Bashir et al., 2015) |
| *IntI1*-R  *IntI1*-F | F29- 305  R29- 530 | AGGAGATCCGAAGACCTC  TCTCGGGTAACATCAAGG | 18  18 | 243 | No | 94^O^–5min; (94^O^–1min; 55^O^–1min; 72^O^–30sec) x35 | Clinical isolates | (Leverstein-Van Hall et al., 2003) |
| Int1A  Int1B | F30- 4  R30- 977 | AAAACCGCCACTGCGCCGTTA  GAAGACGGCTGCACTGAACG | 21  20 | 1201  996*** | No | - | Clinical isolate | (Falbo et al., 1999; Fonseca et al., 2005) |
| Int I.R  Int I.F | F31- 261  R31- 530 | GTTCTTCTACGGCAAGGT  TCTCGGGTAACATCAAGG | 18  18 | 287 | No | 94^O^–5min; (94^O^–20sec; 60^O^–30sec; 72^O^–60sec) x30 | Clinical and animal facility (Faecal samples) | (Kheiri and Akhtari, 2016) |
| F6  R6 | F32- 371  R32- 810 | GCATCCTCGGTTTTCTGG  GGTGTGGCGGGCTTCGTG | 18  18 | 457 | No | 94^O^–2min; (94^O^–1min; 57^O^–1min; 72^O^–90sec) x30 | Clinical (laboratory strains) | (Shibata et al., 2003) |
| *intI*1F  *intI*1R | F33- 111  R33- 797 | TGTCCACTGGGTTCGTGCCT  GCTTCGTGATGCCTGCTTGTT | 20  21 | 707 | No | 94^O^–5min; (94^O^–30sec; 56^O^–30sec; 72^O^–1min) x30; 72^O^–10min | Food (Raw meat samples) | (Zhou et al., 2019) |
| *intI*1L  *intI*1R | F34- 253  R34- 791 | GCCTTGCTGTTCTTCTACGG  GATGCCTGCTTGTTCTACGG | 20  20 | 558 | No | - | Clinical isolates | (Ng et al., 1999) |
| *int*R  *int*F | F35- 409  R35- 534 | GCCCAGCTTCTGTATGGAAC  CCAAGCTCTCGGGTAACATC | 20  20 | 145 | Yes  (SYBR Premix Ex-Taq) | 95^O^–10sec; (95^O^–10sec; 55^O^–10sec; 72^O^–10sec) x40 | Clinical/ Laboratory strains | (Wei et al., 2011) |
| *int*M1‐U  *int*M1‐D | F36- 209  R36- 753 | ACGAGCGCAAGGTTTCGGT  GAAAGGTCTGGTCATACATG | 19  20 | 565 | No | 94^O^–5min; (94^O^–30sec; 52^O^–30sec; 72^O^–2min) x30; 72^O^–7min | Clinical isolates (Faeces, blood, and urine) | (Su et al., 2006) |
| *Int*1-1  *Int*1-2 | F37- 176  R37- 917 | GCGAAGTCGAGGCATTTCTGTC  ATGCGTGTAAATCATCGTCGTAGAGA | 22  26 | 766 | No | - | Clinical strains | (Rodríguez-Martínez et al., 2007) |
| *intI1*F  *intI1*R  *intI1*-Probe | F38- 167  R38- 219  F38 Pb- 185 | TGGGCAGCAGCGAAGTC  TGCGTGGAGACCGAAACC  AGGCATTTCTGTCCTGGCTGGCG | 17  18  23 | 70 | Yes (*Taq*Man Probe) | - | Clinical isolates (Veterinary or food isolates) | (Bugarel et al., 2011) |
| D, intI  A, intI | F39- 168  R39- 991 | GGGCAGCAGCGAAGTCGAGGC  CTACCTCTCACTAGTGAGGGGCGG | 21  24 | 845 | No | 96^O^–1min; (96^O^–30sec ; 58^O^–30sec) x35; 70^O^–60sec; 70^O^–7min | Environmental (Commercial pet turtle eggs and water ponds) | (Díaz et al., 2006) |
| INT1-R  INT1-F | F40- 42  R40- 922 | CAAGGTTCTGGACAGTTGC  TGCGTGTAAATCATCGTCGT | 19  20 | 900 | No | - | Clinical isolates | (Adabi et al., 2009) |
| *intI1*-R  *intI1*-F | F41- 261  R41- 756 | GTTCTTCTACGGCAAGGTG  GCTGAAAGGTCTGGTCATAC | 19  20 | 515 | No | 94^O^–3min; (94^O^–60sec; - ; 72^O^–10min) x30 | Clinical isolates (Hospitalised patients with nosocomial infections) | (D. Wang et al., 2017) |
| *IntI1*F  *IntI1*R | F42- 520  R42- 971 | AAGGATCGGGCCTTGATGTT  CAGCGCATCAAGCGGTGAGC | 20  20 | 471 | No | 94^O^–5min; (94^O^–1min; 55^O^–1min; 72^O^–1min) x30 | Clinical isolates (Blood, pus and urine) | (Pongpech et al., 2008) |
| *intI1*R  *intI1*F | F43- 408  R43- 797 | CGCCCAGCTTCTGTATGG  TTCGTGATGCCTGCTTGTT | 18  19 | 408*** | No | 94^O^–5min; (94^O^–30sec; 51^O^–40sec; 72^O^–40sec) x35; 72^O^–5min | Clinical isolate | (Gu et al., 2008) |
| *IntI1*-f  *IntI1*-r | F44- 70  R44- 571 | ATACGCTACTTGCATTACAG  GCCCGTGCACGCGACAGCTG | 20  20 | 521 | No | 94^O^–5min; (94^O^–30sec; 51^O^–45sec; 72^O^–1-4min) x35; 72^O^–7min | Clinical isolate (ICU patients) | (Zong et al., 2008) |
| *intI1*R  *intI1*F | F45- 641  R45- 869 | ACGCCCTTGAGCGGAAGTATC  GGTTCGAATGTCGTAACCGC | 21  20 | 248 | No | 94^O^–4min; (94^O^–1min; 65^O^–30sec[-1^O^ per cycle]; 70^O^–2min) x10 touchdown cycles; (94^O^–1min; 55^O^–30ec; 70^O^–2min) x24; 70^O^–5min | Clinical (Human faecal samples and pig rectal swabs) | (Phongpaichit et al., 2007) |
| INT1-R  INT1-F  INT1-mF | F46- 1  R46- 534  R46 Pb- 88 | CATGAAAACCGCCACTGC  CCAAGCTCTCGGGTAACATC  GCCTGTTCGGTTCGTAAGCT | 18  20  20 | 553 | No | - | Clinical isolates (including blood, urine and respiratory specimens) | (Hong et al., 2016) |
| Int1 lower  Int1 upper | F47- 10  R47- 888 | GCCACTGCGCCGTTACCACCGC  ATGGCCGAGCAGATCCTGCACG | 22  22 | 900 | No | 94^O^–5min; (94^O^–30sec; 60^O^–40sec; 72^O^–1min) x35; 72^O^–7min | Clinical isolates | (Frank et al., 2007) |
| Int1-1B  Int1-1A | F48- 1  R48- 994 | ATGAAAACCGCCACTGCGCC  TACCTCTCACTAGTGAGGGG | 20  20 | 1012 | No | - | Clinical isolates | (Yan et al., 2006) |
| class1-B  class1-F | F49- 42  R49- 895 | CAAGGTTCTGGACCAGTTGCG  CGGAATGGCCGAGCAGATC | 21  19 | 871 | No | 94^O^–45sec; 5^O^ below melting temperature–60sec; 72^O^–2min | Environmental isolates (Pig farm) | (Sandvang et al., 2002; Spindler et al., 2012) |
| *IntI*-1R  *IntI*-1F | F50- 28  R50- 761 | CCGCTGCGTTCGGTCAAGGT  GGCGCGCTGAAAGGTCTGGT | 20  20 | 753 | No | - | Clinical isolate (Sputum, pus, blood and urine) | (Piyakul et al., 2012) |
| Int I-R  Int I-F | F51- 454  R51- 920 | CAACTGCGGGTCAAGGAT  CGTGTAAATCATCGTCGTAG | 18  20 | 542  486*** | No | - | Clinical isolates | (Yu et al., 2012) |
| Δ*intI1*-F  Δ*intI1*-R | F52- 50  R52- 634 | TGGACCAGTTGCGTGAGC  TCAAGGGCGTCGGGAAG | 18  17 | 601 | No | - | Poultry (Chicken intestinal content and faeces) | (Lai et al., 2013) |
| int_F  int_R | F53- 143**  R53- 419** | CGATGCGTGGAGACCGAAACCTT  GTAACGCGCTTGCTGCTTGGATGC | 23  24 | 301 | No | 95^O^–5min; (95^O^–1min; 58^O^– 1min; 72^O^–5min[+5sec each extension time cycle]) x35; 72^O^–7min | Clinical isolates | (Krauland et al., 2009) |
| *intI1*-R  *intI1*-F | F54- 30  R54- 799 | GCTGCGTTCGGTCAAGGT  GCTTCGTGATGCCTGCTTG | 18  19 | 788 | No | - | Clinical and Food isolates | (Zhao et al., 2018) |
| Int1-F6  Int1-R6 | F55- 185  R55- 810 | AAGCAGACTTGACCTGA  GGTGTGGCGGGCTTCGTG | 17  18 | 457  643*** | No | - | Clinical isolates | (Kainuma et al., 2018) |
| i1219 (r)  i965 (f) | F56- 641  R56- 869 | ACGCCCTTGAGCGGAAGTATC  CCTTCGAATGCTGTAACCGC | 21  20 | 254  248*** | No | 94^O^–4min; (94^O^–1min; 65^O^–30sec[-1^O^ per cycle]; 70^O^–2min) x10 touchdown cycles; (94^O^–1min; 55^O^–30ec; 70^O^–2min) x24; 70^O^–5min | Clinical/Animal isolates | (Ebner et al., 2004) |
| IntΙ-B  IntΙ-F | F57- 166  R57- 736 | TTGGGCAGCAGCGAAGT  TGATGGCGACGCACGAC | 17  17 | 587 | No | 94^O^–5min; (94^O^–40sec; 58^O^–40ec; 72^O^–45sec) x35; 72^O^–5min | Clinical isolates (Sputum) | (Y. Wang et al., 2017) |
| intl1-f  intl1-1014r | F58- 370 **  R58- 994 ** | GCATCCTCGGTTTTCTGG  CTACCTCTCACTAGTGAGGG | 18  20 | 645 | No | - | Clinical isolates | (Kobayashi et al., 2013) |
| intR  intF | **F59**- 399  **R59-** 879 | TCGTTTGTTCGCCCAGC  CCTGCACGGTTCGAATG | 17  17 | 497 | No | 95^O^–5min; (95^O^–45sec; 50^O^–45ec; 72^O^–1min) x30 | Poultry and Swine isolates | (Chuanchuen et al., 2007) |
| int1LF  int1LR | **F60**- 304  **R60**- 439 | CAGGAGATCGGAAGACCT  TTGCAAACCCTCACTGAT | 18  18 | 153*** | Yes  (SYBR Green I) | - | Swine isolates | (Ekkapobyotin et al., 2008) |
| *intI1*F  *intI1*R | **F61**- 587  **R61**- 710 | GTGGATGGCGGCCTGAAGCC  ATTGCCCAGTCGGCAGCG | 20  18 | 141*** | No | 94^O^–5min; (95^O^–1min; 55^O^–30sec; 72^O^–1min) x30; 72^O^–5min | Animal (Foals faecal sample) | (Kennedy et al., 2018) |
| intA  intB | **F62**- 154**  **R62**- 758** | ACAGGGCAAGCTTAGTAAAGCC  CTCGCTAGAACTTTTGGAAA | 22  20 | 625 | No | (95^O^–1min c; 67^O^–1min c; 72^O^–1min) x30; 72^O^–5min | Environmental (Pig slurry and manured agricultural soils) | (Byrne-Bailey et al., 2009) |
| int1-L  int1-F | **F63**- 31  **R63**- 893 | CTGCGTTCGGTCAAGGTTCT  GGAATGGCCGAGCAGATCCT | 20  20 | 882 | No | **-** | Food isolates (commercial fish and seafood) | (Ryu et al., 2012) |
| *Int1*F  *Int1*R | **F64**- 584  **R64**- 833 | CACGGATATGCGACAAAAAG  GATGACAACGAGTGACGAAATG | 20  22 | 160  271*** | No | 94^O^–5min; (94^O^–1min; 51^O^–1min; 72^O^–45sec) x35; 72^O^–5min | Clinical (Blood) and Environmental (Tap water) | (Karami et al., 2020) |

F, forward primer; R, reverse primer; P, probe

* denotes hit start position of primer based on alignment to a reference *intI1* gene sequence shown in Figure S.1 ** denotes hit position based on Primer prospector alignment of CP003684.1 *** denotes estimated amplicon size based on alignment of primer to CP003684.1 *intI*1 reference sequence**.**

**Table S.2** Listed of *intI*1 primer sets excluded from further analysis in this study

F, forward primer; R, reverse primer; W.S, weighted score. A, start position of primers on intI1 sequence based on alignment to the reference *Pseudomonas aeruginosa* plasmid pVS1 nucleotide sequence (Figure S.1). b, start position of primers on *intI*1 sequence based on alignment with Primer Prospector to sequence within the complete length intI1 sub-database, SDB1 (*n=104*). Mean WS

| Primer Pair | Sequence (5’-3’ Direction) | Position^a^ | Position^b^ | Mean W.S* | Expected amplicon size (bp) | Observed amplicon size (bp) | Comment(s) | Study reference |
| --- | --- | --- | --- | --- | --- | --- | --- | --- |
| F11  R11 | CTTCAGCCTTTTCCAGCAAC  GAAACCTGCTCCAGCACTTC | 158  956 | 157  42 | 3.41  6.42 | 308 | 818 | Large difference between observed and expected amplicon size coupled to a high mean weighted score for the primers | (Najafgholizadeh Pirzaman and Mojtahedi, 2019) |
| F12  R12 | TCATGGCTTGTTATGACTGT  GTAGGGCTTATTATGCACGC | 973  752 | 972  886 | 7.22  5.76 | 600 | - | Primer sequence same as those of primer set hep58 and hep59, commonly used to target cassette region of class 1 integron. | (Mobaraki et al., 2018) |
| F20  R20 | AGCTTACGAACCGAACAGGC  TCCGCCAGGATTGACTTGCG | 88  666 | 87  665 | 0.37  3.69 | 950 | 597 | Large difference between estimated and expected amplicon size | (Borruso et al., 2016) |
| F24  R24 | CCCGAGGCATAGACTGTA  CAGTGGACATAAGCCTGTTC | 238  100 | 227  99 | 6.88  0.15 | 160 | - | Wrong primer binding orientation even when the forward sequence is used as the reverse sequence and vice versa | (Koeleman et al., 2001) |
| F27  R27 | CGAGGCATAGACTGTAC  TTCGAATGTCGTAACCGC | 876  869 | 182  868 | 2.81  0.01 | 925 | 703 | Large difference between observed and expected amplicon size | (Orman et al., 2002) |
| F28  R28 | GTCAAGGTTCTGGACCAGTTG  ATCATCGTCGTAGAGACGTCGG | 40  911 | 39  910 | **5.4**  **4.64** | 550 | 892 | Large difference between estimated and expected amplicon size; primer sequence same as F17-R17 (Rosser and Young, 1999) with 1bp difference in the forward sequence. | (Bashir et al., 2015) |
| F30  R30 | AAAACCGCCACTGCGCCGTTA  GAAGACGGCTGCACTGAACG | 4  977 | 3  275 | 0.19  6.25 | 1201 | 996 | Large difference between expected and observed amplicon size. Additionally, expected amplicon size exceeds the size of a complete length *intI*1 gene (1014bp) | (Falbo et al., 1999; Fonseca et al., 2005) |
| F55  R55 | AAGCAGACTTGACCTGA  GGTGTGGCGGGCTTCGTG | 185  810 | 184  809 | 5.46  0.09 | 457 | 643 | Large difference between observed and expected amplicon size | (Kainuma et al., 2018) |
| F61  R61 | GTGGATGGCGGCCTGAAGCC  ATTGCCCAGTCGGCAGCG | 587  710 | 586  319 | 4.41  3.99 | - | 141 | Targets aminoglycoside adenylyl transferases (*aadA1a;* previously *ant(3”)Ia*) gene (Sandvang et al., 1997; Guerra et al., 2001) | (Kennedy et al., 2018) |
| F64  R64 | CACGGATATGCGACAAAAAG  GATGACAACGAGTGACGAAATG | 584  833 | 935  832 | 5.02  4.99 | 160 | 271 | Targets Class 2 integron-integrase (*intI*2) gene (Gündoǧdu et al., 2011) | (Karami et al., 2020) |

**Table S.****3:** Coverage and specificity of currently published and newly modified *intI*1 primer pairs

| **PCR Primer Set** | **Coverage Test** | | | | | | | | | **Specificity Test** | | | | | |
| --- | --- | --- | --- | --- | --- | --- | --- | --- | --- | --- | --- | --- | --- | --- | --- |
|  | ***intI1* Sub-databases** | | | | | | | | | ***intI1-Like sequence (n= 15)*** | | | ***Non-intI1 sequence (n= 1540)*** | | |
|  | **SDB1**  ***(n= 104)*** | | | **SDB2 *(n= 144)*** | | | **SDB3 *(n=503)*** | | |  |  |  |  |  |  |
|  | **Number of sequences with correct primer orientation (%)** | **Mean Weighted Score** | **Target amplified (%) (0 Mismatches)** | **Number of sequences with correct primer orientation (%)** | **Mean Weighted Score** | **Target amplified (%) (0 mismatch)** | **Number of sequences with correct primer orientation (%)** | **Mean Weighted Score** | **Target amplified (%) (0 mismatch)** | **Number of sequences with correct primer orientation (%)** | **Mean Weighted Score** | **Target amplified (%) (0 mismatch)** | **Number of sequences with correct primer orientation (%)** | **Mean Weighted Score** | **Target amplified (%) (0 mismatch)** |
| **DF-DR** | 104 (100%) | **F** 0.008 | 102 (98%) | 144 (100%) | **F** 0.006 | 142 (99%) | 502 (100%) | **F** 0.016 | 493 (99%) | 15 (100%) | **F** 0.6 | 9 (60%) | 813 (53%) | **F** 4.991 | 0 (0%) |
|  |  | **R** 0 |  |  | **R** 0 |  |  | **R** 0.002 |  |  | **R** 0.07 |  |  | **R** 4.223 |  |
| **F1-R1** | 104 (100%) | **F** 0.017 | 101 (97%) | 144 (100%) | **F** 0.013 | 141 (98%) | 458 (91%) | **F** 0.041 | 448 (98%) | 14 (93%) | **F** 0.4 | 9  (64%) | 736 (48%) | **F** 4.342 | 0 (0%) |
|  |  | **R** 0.004 |  |  | **R** 0.003 |  |  | **R** 0.002 |  |  | **R** 0.057 |  |  | **R** 4.493 |  |
| **F2-R2** | 104 (100%) | **F** 0.019 | 97 (93%) | 143 (99%) | **F** 0.014 | 133 (93%) | 443 (88%) | **F** 0.009 | 413 (94%) | 11 (73 %) | **F** 0.491 | 7 (64%) | 923 (60%) | **F** 5.215 | 0 (0%) |
|  |  | **R** 0.042 |  |  | **R** 0.073 |  |  | **R** 0.08 |  |  | **R** 0.782 |  |  | **R** 5.992 |  |
| **F3-R3** | 104 (100%) | **F** 0.012 | 101 (97%) | 144 (100%) | **F** 0.008 | 141 (98%) | 501 (100%) | **F** 0.003 | 490 (98%) | 14 (93%) | **F** 0.171 | 8 (57%) | 997 (65%) | **F** 6.024 | 0 (0%) |
|  |  | **R** 0 |  |  | **R** 0 |  |  | **R** 0.006 |  |  | **R** 0.314 |  |  | **R** 5.579 |  |
| **F4-R4** | 104 (100%) | **F** 0.037 | 98 (94%) | 144 (100%) | **F** 0.026 | 138  (96%) | 441 (88%) | **F** 0.034 | 428 (97%) | 14 (93%) | **F** 0.671 | 9 (64%) | 879 (57%) | **F** 5.94 | 0 (0%) |
|  |  | **R** 0.05 |  |  | **R** 0.036 |  |  | **R** 0.044 |  |  | **R** 0.529 |  |  | **R** 5.534 |  |
| **F5-R5** | 104 (100%) | **F** 0.079 | 96 (92%) | 140 (97%) | **F** 0.059 | 118 (84%) | 455 (90%) | **F** 0.596 | 315 (70%) | 14 (93%) | **F** 2.371 | 6 (43%) | 912 (59%) | **F** 4.389 | 0 (0%) |
|  |  | **R** 0.035 |  |  | **R** 0.169 |  |  | **R** 1.026 |  |  | **R** 2.929 |  |  | **R** 4.968 |  |
| **F6-R6** | 104 (100%) | **F** 0.019 | 94 (90%) | 127 (88%) | **F** 0.052 | 116 (91%) | 318 (63%) | **F** 0.028 | 294 (93%) | 9  (60%) | **F** 0.089 | 7 (78%) | 972 (63%) | **F** 5.37 | 0 (0%) |
|  |  | **R** 0.06 |  |  | **R** 0.085 |  |  | **R** 0.145 |  |  | **R** 0.267 |  |  | **R** 4.246 |  |
| **F7-R7** | 104 (100%) | **F** 0.046 | 101 (97%) | 144 (100%) | **F** 0.065 | 140 (97%) | 475 (94%) | **F** 0.02 | 469 (99%) | 10 (67%) | **F** 0.260 | 8 (80%) | 778 (51%) | **F** 5.882 | 0 (0%) |
|  |  | **R** 0.008 |  |  | **R** 0.031 |  |  | **R** 0.009 |  |  | **R** 0.080 |  |  | **R** 4.636 |  |
| **F8-R8** | 101 (97%) | **F** 0 | 101 (100%) | 115 (80%) | **F** 0 | 114 (99%) | 243 (48%) | **F** 0.005 | 221 (92%) | 8  (53%) | **F** 0.05 | 5 (63%) | 1194 (78%) | **F** 4.617 | 0  (0%) |
|  |  | **R** 0 |  |  | **R** 0.012 |  |  | **R** 0.113 |  |  | **R** 0.525 |  |  | **R** 4.634 |  |
| **F9-R9** | 104 (100%) | **F** 0.396 | 0 (0%) | 141 (98%) | **F** 0.372 | 0 (0%) | 420 (83%) | **F** 0.345 | 0 (0%) | 13 (87%) | **F** 0.615 | 0 (0%) | 848 (55%) | **F** 5.302 | 0  (0%) |
|  |  | **R** 5.019 |  |  | **R** 5.014 |  |  | **R** 5.005 |  |  | **R** 5.108 |  |  | **R** 5.055 |  |
| **F10-R10** | 104 (100%) | **F** 0.063 | 93 (89%) | 126 (88%) | **F** 0.052 | 115 (91%) | 302 (60%) | **F** 0.022 | 289 (96%) | 9  (60%) | **F** 1.178 | 6 (67%) | 908 (59%) | **F** 5.438 | 0 (0%) |
|  |  | **R** 0.117 |  |  | **R** 0.097 |  |  | **R** 0.044 |  |  | **R** 0.8 |  |  | **R** 5.549 |  |
| **F13-R13** | 104 (100%) | **F** 0.013 | 101 (97%) | 144 (100%) | **F** 0.01 | 139 (97%) | 473 (94%) | **F** 0.392 | 431 (92%) | 13 (87%) | **F** 0.492 | 9 (69%) | 870 (56%) | **F** 4.226 | 0 (0%) |
|  |  | **R** 0.004 |  |  | **R** 0.008 |  |  | **R** 0.231 |  |  | **R** 0.338 |  |  | **R** 4.85 |  |
| **F14-R14** | 102 (98%) | **F** 0 | 101 (99%) | 117 (81%) | **F** 0 | 114 (97%) | 227 (45%) | **F** 0.004 | 217 (96%) | 6  (40%) | **F** 0.133 | 4 (67%) | 1055 (69%) | **F** 4.052 | 14 (0.9%) |
|  |  | **R** 0.025 |  |  | **R** 0.055 |  |  | **R** 0.041 |  |  | **R** 0.333 |  |  | **R** 2.956 |  |
| **F15-R15** | 101 (97%) | **F** 0 | 101 (100%) | 115 (80%) | **F** 0 | 114 (99%) | 243 (48%) | **F** 0.002 | 223 (93%) | 8  (53%) | **F** 0.05 | 5 (63%) | 1051 (68%) | F 5.496 | 0 (0%) |
|  |  | **R** 0 |  |  | **R** 0.012 |  |  | **R** 0.113 |  |  | **R** 0.525 |  |  | R 4.356 |  |
| **F16-R16** | 104 (100%) | **F** 0.008 | 102 (98%) | 144 (100%) | **F** 0.019 | 141 (98%) | 427 (85%) | **F** 0.02 | 415 (98%) | 14 (93%) | **F** 0.3 | 11 (79%) | 961 (62%) | **F** 5.64 | 0 (0%) |
|  |  | **R** 0 |  |  | **R** 0 |  |  | **R** 0.014 |  |  | **R** 0.029 |  |  | **R** 5.543 |  |
| **F17-R17** | 104 (100%) | **F** 0.067 | 98 (94%) | 142 (99%) | **F** 0.444 | 104 (73%) | 466 (93%) | **F** 1.028 | 154 (33%) | 15 (100%) | **F** 0.827 | 3 (20%) | 787 (51%) | **F** 5.847 | 0 (0%) |
|  |  | **R** 0.121 |  |  | **R** 0.866 |  |  | **R** 2.596 |  |  | **R** 2.360 |  |  | **R** 5.709 |  |
| **F18-R18** | 104 (100%) | **F** 0.04 | 98 (94%) | 144 (100%) | **F** 0.246 | 100 (69%) | 484 (96%) | **F** 0.568 | 128 (27%) | 14 (93%) | **F** 0.257 | 2 (14%) | 814 (53%) | **F** 4.153 | 0 (0%) |
|  |  | **R** 0.121 |  |  | **R** 1.158 |  |  | **R** 2.632 |  |  | **R** 3.186 |  |  | **R** 4.17 |  |
| **F19-R19** | 104 (100%) | **F** 0.035 | 98 (94%) | 142 (99%) | **F** 0.166 | 108 (76%) | 482 (96%) | **F** 0.438 | 168 (35%) | 15 (100%) | **F** 0.347 | 4 (27%) | 711 (46%) | **F** 4.44 | 0 (0%) |
|  |  | **R** 0.115 |  |  | **R** 0.465 |  |  | **R** 1.413 |  |  | **R** 1.280 |  |  | **R** 4.224 |  |
| **F21-R21** | 104 (100%) | **F** 0.004 | 99 (95%) | 144 (100%) | **F** 0.003 | 121 (84%) | 501 (99%) | **F** 0.427 | 272 (55%) | 14 (93%) | **F** 0.629 | 4 (29%) | 800 (52%) | **F** 4.659 | 0 (0%) |
|  |  | **R** 0.048 |  |  | **R** 0.542 |  |  | **R** 1.554 |  |  | **R** 1.957 |  |  | **R** 4.509 |  |
| **F22-R22** | 104 (100%) | **F** 0.015 | 97 (93%) | 144 (100%) | **F** 0.025 | 134 (93%) | 462 (92%) | **F** 0.879 | 377 (82%) | 10 (71%) | **F** 2.020 | 5 (50%) | 872 (57%) | **F** 4.962 | 0 (0%) |
|  |  | **R** 0.062 |  |  | **R** 0.075 |  |  | **R** 0.06 |  |  | **R** 0.08 |  |  | **R** 5.87 |  |
| **F23-R23** | 101 (97%) | **F** 0 | 101 (100%) | 115 (80%) | **F** 0 | 114 (99%) | 245 (49%) | **F** 0.033 | 221 (91%) | 8  (53%) | **F** 0.05 | 5 (63%) | 1149 (75%) | **F** 4.559 | 0 (0%) |
|  |  | **R** 0 |  |  | **R** 0.012 |  |  | **R** 0.17 |  |  | **R** 0.525 |  |  | **R** 5.013 |  |
| **F25-R25** | 102 (98%) | **F** 0.096 | 91 (89%) | 130 (90%) | **F** 0.075 | 99 (76%) | 381 (76%) | **F** 0.13 | 162 (43%) | 13 (87%) | **F** 0.446 | 4 (31%) | 843 (55%) | **F** 5.735 | 0 (0%) |
|  |  | **R** 0.041 |  |  | **R** 0.755 |  |  | **R** 2.696 |  |  | **R** 2.492 |  |  | **R** 5.161 |  |
| **F26-R26** | 102 (98%) | **F** 0.096 | 95 (93%) | 130 (90%) | **F** 0.075 | 121 (93%) | 380 (76%) | **F** 0.111 | 361 (96%) | 13 (87%) | **F** 0.446 | 8 (62%) | 659 (43%) | **F** 5.926 | 0 (0%) |
|  |  | **R** 0.008 |  |  | **R** 0.012 |  |  | **R** 0.084 |  |  | **R** 0.138 |  |  | **R** 5.661 |  |
| **F29-R29** | 104 (100%) | **F** 0.408 | 0  (0%) | 143 (99%) | **F** 0.406 | 0 (0%) | 489 (97%) | **F** 0.442 | 0 (0%) | 13 (87%) | **F** 0.462 | 0 (0%) | 661 (43%) | **F** 4.557 | 0 (0%) |
|  |  | **R** 0.017 |  |  | **R** 0.013 |  |  | **R** 0.004 |  |  | **R** 0.431 |  |  | **R** 4.826 |  |
| **F31-R31** | 104 (100%) | **F** 0.004 | 101 (97%) | 143 (99%) | **F** 0.027 | 138 (97%) | 502 (100%) | **F** 0.263 | 464 (93%) | 14 (93%) | **F** 0.514 | 8 (57%) | 674 (44%) | **F** 4.769 | 0 (0%) |
|  |  | **R** 0.017 |  |  | **R** 0.013 |  |  | **R** 0.004 |  |  | **R** 0.4 |  |  | **R** 4.572 |  |
| **F32-R32** | 104 (100%) | **F** 0.012 | 97 (93%) | 126 (88%) | **F** 0.01 | 119 (94%) | 324 (64%) | **F** 0.085 | 294 (91%) | 8  (53%) | **F** 0.650 | 4 (50%) | 896 (58%) | **F** 4.929 | 0 (0%) |
|  |  | **R** 0.075 |  |  | **R** 0.062 |  |  | **R** 0.223 |  |  | **R** 0.45 |  |  | **R** 4.569 |  |
| **F33-R33** | 104 (100%) | **F** 0.098 | 78 (75%) | 137 (95%) | **F** 0.088 | 97 (71%) | 432 (86%) | **F** 0.606 | 196 (46%) | 13 (87%) | **F** 0.754 | 2 (15%) | 903 (59%) | **F** 5.245 | 0 (0%) |
|  |  | **R** 0.06 |  |  | **R** 0.693 |  |  | **R** 2.308 |  |  | **R** 2.923 |  |  | **R** 5.266 |  |
| **F34-R34** | 104 (100%) | **F** 0.033 | 98 (94%) | 144 (100%) | **F** 0.031 | 120 (83%) | 502 (100%) | **F** 0.324 | 272 (54%) | 14 (93%) | **F** 0.471 | 4 (29%) | 787 (51%) | **F** 5.511 | 0  (0%) |
|  |  | **R** 0.042 |  |  | **R** 0.771 |  |  | **R** 2.363 |  |  | **R** 2.371 |  |  | **R** 5.18 |  |
| **F35-R35** | 104 (100%) | **F** 0 | 102 (98%) | 143 (99%) | **F** 0.003 | 140 (98%) | 502 (100%) | **F** 0.012 | 493 (99%) | 14 (93%) | **F** 0. 3 | 10 (71%) | 808 (52%) | **F** 5.52 | 0 (0%) |
|  |  | **R** 0.015 |  |  | **R** 0.011 |  |  | **R** 0.003 |  |  | **R** 0.186 |  |  | **R** 5.068 |  |
| **F36-R36** | 104 (100%) | **F** 0.069 | 93 (89%) | 143 (99%) | **F** 0.067 | 124 (87%) | 441 (88%) | **F** 0.771 | 349 (79%) | 10 (67%) | **F** 1.62 | 5 (50%) | 703 (46%) | **F** 5.239 | 0 (0%) |
|  |  | **R** 0.033 |  |  | **R** 0.057 |  |  | **R** 0.066 |  |  | **R** 1.28 |  |  | **R** 5.468 |  |
| **F37-R37** | 104 (100%) | **F** 0.037 | 98 (94%) | 144 (100%) | **F** 0.026 | 111 (77%) | 502 (100%) | **F** 0.619 | 167 (33%) | 15 (100%) | **F** 1.107 | 2 (13%) | 1105 (72%) | **F** 5.73 | 0 (0%) |
|  |  | **R** 0.148 |  |  | **R** 1.11 |  |  | **R** 2.888 |  |  | **R** 2.88 |  |  | **R** 6.227 |  |
| **F38-R38** | 104 (100%) | **F** 0 | 95 (91%) | 144 (100%) | **F** 0 | 132 (92%) | 438 (87%) | **F** 0.017 | 419 (96%) | 13 (87%) | **F** 0.262 | 9  (69%) | 801 (52%) | **F** 4.829 | 0 (0%) |
|  |  | **R** 0.083 |  |  | **R** 0.081 |  |  | **R** 0.042 |  |  | **R** 0.185 |  |  | **R** 4.502 |  |
| **F39-R39** | 104 (100%) | **F** 0.01 | 76 (73%) | 144 (100%) | **F** 0.007 | 87 (60%) | 493 (98%) | **F** 0.875 | 107 (22%) | 13 (87%) | **F** 1.138 | 1 (8%) | 751 (49%) | **F** 5.503 | 0 (0%) |
|  |  | **R** 0.919 |  |  | **R** 1.714 |  |  | **R** 3.318 |  |  | **R** 4.985 |  |  | **R** 6.96 |  |
| **F40-R40** | 104 (100%) | **F** 1.033 | 1 (0.9%) | 142 (99%) | **F** 1.165 | 1 (0.7%) | 466 (93%) | **F** 1.389 | 1 (0.2%) | 15 (100%) | **F** 1.293 | 0 (0%) | 1146 (74%) | **F** 4.994 | 0 (0%) |
|  |  | **R** 0.165 |  |  | **R** 0.987 |  |  | **R** 2.593 |  |  | **R** 2.653 |  |  | **R** 3.732 |  |
| **F41-R41** | 104 (100%) | **F** 0.004 | 102 (98%) | 143 (99%) | **F** 0.013 | 136 (95%) | 442 (88%) | **F** 0.511 | 389 (88%) | 11 (73 %) | **F** 0.873 | 5 (45%) | 699 (45%) | **F** 4.971 | 0 (0%) |
|  |  | **R** 0.004 |  |  | **R** 0.062 |  |  | **R** 0.1 |  |  | **R** 0.509 |  |  | **R** 4.945 |  |
| **F42-R42** | 101 (97%) | **F** 0.004 | 90 (89%) | 114 (79%) | **F** 0.04 | 102 (89%) | 208 (41%) | **F** 0.024 | 190 (92%) | 4  (27%) | **F** 0.2 | 2 (50%) | 757 (49%) | **F** 4.577 | 0 (0%) |
|  |  | **R** 0.17 |  |  | **R** 0.195 |  |  | **R** 0.138 |  |  | **R** 1.750 |  |  | **R** 4.857 |  |
| **F43-R43** | 104 (100%) | **F** 0 | 99 (95%) | 137 (95%) | **F** 0.003 | 121 (88%) | 439 (87%) | **F** 0.005 | 300 (69%) | 11 (73 %) | **F** 0 | 6 (55%) | 748 (49%) | **F** 4.74 | 0 (0%) |
|  |  | **R** 0.044 |  |  | **R** 0.472 |  |  | **R** 1.604 |  |  | **R** 1.964 |  |  | **R** 4.518 |  |
| **F44-R44** | 104 (100%) | **F** 0.06 | 97 (93%) | 139 (97%) | **F** 0.045 | 131 (94%) | 403 (80%) | **F** 0.041 | 387 (97%) | 13 (87%) | **F** 0.138 | 6 (46%) | 835 (54%) | **F** 5.151 | 0 (0%) |
|  |  | **R** 0.012 |  |  | **R** 0.035 |  |  | **R** 0.015 |  |  | **R** 0.692 |  |  | **R** 5.451 |  |
| **F45-R45** | 104 (100%) | **F** 0.015 | 100 (96%) | 126 (88%) | **F** 0.056 | 120 (95%) | 316 (63%) | **F** 0.023 | 281 (89%) | 9  (60%) | **F** 0.044 | 8 (89%) | 947 (61%) | **F** 5.508 | 0 (0%) |
|  |  | **R** 0 |  |  | **R** 0.037 |  |  | **R** 0.409 |  |  | **R** 0.111 |  |  | **R** 4.907 |  |
| **F46-R46** | 102 (98%) | **F** 0.041 | 95 (93%) | 131 (91%) | **F** 0.079 | 121 (92%) | 386 (77%) | **F** 0.166 | 363 (95%) | 13 (87%) | **F** 0.246 | 8 (62%) | 767 (50%) | **F** 4.834 | 0 (0%) |
|  |  | **R** 0.016 |  |  | **R** 0.069 |  |  | **R** 0.023 |  |  | R 0.677 |  |  | **R** 4.968 |  |
| **F47-R47** | 102 (98%) | **F** 0.057 | 91 (89%) | 130 (90%) | **F** 0.045 | 99 (76%) | 382 (76%) | **F** 0.114 | 161 (42%) | 13 (87%) | **F** 0.446 | 4 (31%) | 707 (46%) | **F** 5.875 | 0 (0%) |
|  |  | **R** 0.057 |  |  | **R** 0.808 |  |  | **R** 2.895 |  |  | **R** 2.677 |  |  | **R** 5.318 |  |
| **F48-R48** | 103 (99%) | **F** 0.134 | 71 (69%) | 143 (99%) | **F** 0.55 | 72 (50%) | 486 (97%) | **F** 1.209 | 77 (16%) | 13 (87%) | **F** 0.554 | 2 (15%) | 880 (57%) | **F** 5.306 | 0 (0%) |
|  |  | **R** 0.746 |  |  | **R** 1.432 |  |  | **R** 2.807 |  |  | **R** 4.523 |  |  | **R** 5.722 |  |
| **F49-R49** | 104 (100%) | **F** 0.081 | 95 (91%) | 142 (99%) | **F** 0.552 | 103 (73%) | 468 (93%) | **F** 1.309 | 167 (36%) | 15 (100%) | **F** 1.053 | 5 (33%) | 839 (54%) | **F** 5.465 | 0 (0%) |
|  |  | **R** 0.075 |  |  | **R** 0.449 |  |  | **R** 1.229 |  |  | **R** 1.24 |  |  | **R** 4.357 |  |
| **F50-R50** | 104 (100%) | **F** 0.081 | 101 (97%) | 144 (100%) | **F** 0.59 | 125 (87%) | 493 (98%) | **F** 1.281 | 301 (61%) | 10 (67%) | **F** 0.58 | 7 (70%) | 782 (51%) | **F** 5.203 | 0 (0%) |
|  |  | **R** 0.008 |  |  | **R** 0.124 |  |  | **R** 0.534 |  |  | **R** 1.0 |  |  | **R** 5.264 |  |
| **F51-R51** | 102 (97%) | **F** 0 | 101 (99%) | 118 (82%) | **F** 0 | 114 (97%) | 256 (51%) | **F** 0.017 | 227 (89%) | 8  (53%) | **F** 0.15 | 5 (63%) | 807 (52%) | **F** 4.188 | 0 (0%) |
|  |  | **R** 0.029 |  |  | **R** 0.097 |  |  | **R** 0.184 |  |  | **R** 0.35 |  |  | **R** 5.266 |  |
| **F52-R52** | 104 (100%) | **F** 0.081 | 98 (94%) | 144 (100%) | **F** 0.392 | 125 (87%) | 503 (100%) | **F** 0.95 | 369 (74%) | 15 (100%) | **F** 0.52 | 9 (60%) | 823 (53%) | **F** 4.126 | 0  (0%) |
|  |  | **R** 0.008 |  |  | **R** 0.026 |  |  | **R** 0.061 |  |  | **R** 1.32 |  |  | **R** 4.243 |  |
| **F53-R53** | 104 (100%) | **F** 4.219 | 0 (0%) | 144 (100%) | **F** 4.214 | 0 (0%) | 426 (85%) | **F** 4.208 | 0 (0%) | 13 (87%) | **F** 4.231 | 0 (0%) | 715 (46%) | **F** 5.867 | 0 (0%) |
|  |  | **R** 5.65 |  |  | **R** 5.631 |  |  | **R** 5.612 |  |  | **R** 6.4 |  |  | **R** 6.198 |  |
| **F54-R54** | 104 (100%) | **F** 0.079 | 97 (93%) | 142 (99%) | **F** 0 | 121 (85%) | 475 (94%) | **F** 1.096 | 211 (45%) | 14 (93%) | **F** 0.357 | 5 (36%) | 893 (60%) | **F** 4.744 | 0 (0%) |
|  |  | **R** 0.06 |  |  | **R** 0.748 |  |  | **R** 2.215 |  |  | **R** 3.171 |  |  | **R** 5.006 |  |
| **F56-R56** | 104 (100%) | **F** 0.015 | 0 (0%) | 141 (98%) | **F** 0.011 | 0 (0%) | 453 (90%) | **F** 0.004 | 0 (0%) | 10 (67%) | **F** 0.04 | 0 (0%) | 674 (44%) | **F** 5.644 | 0 (0%) |
|  |  | **R** 1.6 |  |  | **R** 1.923 |  |  | **R** 2.620 |  |  | **R** 1.98 |  |  | **R** 4.724 |  |
| **F57-R57** | 103 (99%) | **F** 0 | 101 (98%) | 142 (99%) | **F** 0 | 140 (99%) | 459 (91%) | **F** 0.458 | 381 (83%) | 12 (80%) | **F** 0.667 | 7 (58%) | 636 (41%) | **F** 4.115 | 0 (0%) |
|  |  | **R** 0.008 |  |  | **R** 0.006 |  |  | **R** 0.014 |  |  | **R** 0.533 |  |  | **R** 4.273 |  |
| **F58-R58** | 104 (100%) | **F** 0.012 | 75 (72%) | 144 (100%) | **F** 0.008 | 86 (60%) | 503 (100%) | **F** 0.055 | 158 (32%) | 13 (87%) | **F** 0.4 | 1 (8%) | 943 (61%) | **F** 4.866 | 0 (0%) |
|  |  | **R** 0.812 |  |  | **R** 1.497 |  |  | **R** 2.937 |  |  | **R** 4.523 |  |  | **R** 5.583 |  |
| **F59-R59** | 104 (100%) | **F** 0 | 103 (99%) | 124 (86%) | **F** 0 | 123 (99%) | 314 (62%) | **F** 0.012 | 277 (88%) | 9  (60%) | **F** 0.378 | 8 (89%) | 1082 (70%) | **F** 4.613 | 0 (0%) |
|  |  | **R** 0.004 |  |  | **R** 0.003 |  |  | **R** 0.206 |  |  | **R** 0 |  |  | **R** 3.979 |  |
| **F60-R60** | 104 (100%) | F 0.008 | 101 (97%) | 144 (100%) | **F** 0.006 | 141  (98%) | 487 (97%) | **F** 0.002 | 482 (99%) | 14 (93%) | **F** 0.086 | 10 (71%) | 774 (50%) | **F** 5.16 | 0 (0%) |
|  |  | R 0.04 |  |  | **R** 0.003 |  |  | **R** 0.001 |  |  | **R** 0.214 |  |  | **R** 4.302 |  |
| **F62-R62** | 102 (98%) | **F** 8.233 | 0 (0%) | 141 (98%) | **F** 8.241 | 0  (0%) | 437 (87%) | **F** 8.174 | 0 (0%) | 13 (87%) | **F** 7.831 | 0 (0%) | 638 (41%) | **F** 5.617 | 0 (0%) |
|  |  | **R** 4.418 |  |  | **R** 4.472 |  |  | **R** 4.717 |  |  | **R** 5.354 |  |  | **R** 4.934 |  |
| **F63-R63** | 104 (100%) | **F** 0.056 | 96 (92%) | 142 (99%) | **F** 0.28 | 104 (73%) | 471  (94%) | **F** 0.663 | 170 (36%) | 15 (100%) | **F** 0.52 | 6 (40%) | 815 (53%) | **F** 5.542 | 0 (0%) |
|  |  | **R** 0.069 |  |  | **R** 0.794 |  |  | **R** 2.479 |  |  | **R** 2.36 |  |  | **R** 4.22 |  |

F, forward primer; R, reverse primer; SDB, sub-database. WS is the sum of Weighted score is the sum of score for the forward and reverse primer for each primer set.

**Table S** **4:** intI1 primer sets with no amplicon produced at 0 WS and the WS at which an amplicon was produced

| Primer pair | SDB1 *(n=104)* | | SDB2 *(n=144)* | | SDB3 *(n=503)* | |
| --- | --- | --- | --- | --- | --- | --- |
|  | **WS at which an amplicon was produced** | **No of amplicon generated** | **WS at which an amplicon was produced** | **No of amplicon generated** | **WS at which an amplicon was produced** | **No of amplicon generated** |
| **F9-R9** | 5 | 17 | 5 | 27 | 5 | 109 |
| **F29-R29** | 0.4 | 100 | 0.4 | 139 | 0.4 | 482 |
| **F53-R53** | 9.2 | 1 | 9 | 1 | 9 | 1 |
| **F56-R56** | 1.6 | 100 | 1.6 | 120 | 0.8 | 1 |
| **F62-R62** | 8.7 | 1 | 8.7 | 1 | 6.2 | 1 |

F, forward primer; R, reverse primer; SDB, sub-database; WS, Weighted score

| **Primer pair and probe combination** | **SDB1 *(n= 104)*** | | | | **SDB2**  ***(n= 144)*** | | | | **SDB3 *(n= 503)*** | | | | ***intI1*-like**  ***(n=15)*** | | | |
| --- | --- | --- | --- | --- | --- | --- | --- | --- | --- | --- | --- | --- | --- | --- | --- | --- |
|  | **Number of sequences with correct primer orientation (%)** | **Weighted Score Threshold** | | | **Number of sequences with correct primer orientation (%)** | **Weighted Score Threshold** | | | **Number of sequences with correct primer orientation (%)** | **Weighted Score Threshold** | | | **Number of sequences with correct primer orientation (%)** | **Weighted Score Threshold** | | |
|  |  | **0 (0 mismatch)** | **0.4 ( 1 mismatch at 5’end)** | **1 (mismatch at 3’end/non- 3’gaps/ 2 mismatches at 5’end)** |  | **0 (0 mismatch)** | **0.4 ( 1 mismatch at 5’end)** | **1 (mismatch at 3’end/non- 3’gaps/ 2 mismatches at 5’end)** |  | **0 (0 mismatch)** | **0.4 ( 1 mismatch at 5’end)** | **1 (mismatch at 3’end/non- 3’gaps/ 2 mismatches at 5’end)** |  | **0 (0 mismatch)** | **0.4 ( 1 mismatch at 5’end)** | **1 (mismatch at 3’end/non- 3’gaps/ 2 mismatches at 5’end)** |
| **DF-P-DR** | 104 (100%) | 102 (98%) | 104 (100%) | 104 (100%) | 144 (100%) | 142 (99%) | 144 (100%) | 144 (100%) | 501 (100%) | 494 (99%) | 501 (100%) | 501 (100%) | 14 (93%) | 9  (64%) | 11 (79%) | 12 (86%) |
| **F7-P-R7** | 104 (100%) | 92 (88%) | 96 (92%) | 102 (98%) | 144 (100%) | 131 (91%) | 135 (94%) | 141 (98%) | 475 (94%) | 454 (96%) | 465 (98%) | 471 (99%) | 10 (67%) | 8  (80%) | 8  (80%) | 8  (80%) |
| **F10-P-R10** | 104 (100%) | 91 (88%) | 92 (88%) | 99 (95%) | 126 (88%) | 113 (90%) | 114 (90%) | 121 (96%) | 302 (60%) | 288 (95%) | 289 (96%) | 297 (98%) | 9  (60%) | 6  (67%) | 6  (67%) | 6  (67%) |
| **F38-P-R38** | 104 (100%) | 92 (88%) | 96 (92%) | 103 (99%) | 144 (100%) | 127 (88%) | 131 (91%) | 143 (99%) | 438 (87%) | 415 (95%) | 419 (96%) | 435 (99%) | 13 (87%) | 9  (69%) | 9  (69%) | 9  (69%) |
| **F46-P-R46** | 102  (98%) | 34 (33%) | 54 (53%) | 95 (93%) | 130 (90%) | 41 (32%) | 67 (52%) | 121 (93%) | 380 (76%) | 88 (23%) | 185 (49%) | 360 (95%) | 13 (87%) | 3  (23%) | 3  (23%) | 8  (23%) |

**Table S.****5:** Coverage of published and newly modified intI1 primer sets that incorporates a reporter probe

F, forward primer; R, reverse primer; P, probe; SDB, sub-database

| **TukeyHSD** | ***P-*adjusted value** |
| --- | --- |
| CST-Household Influent-F3-R3 : CST-Household Influent-DF-DR | 0.981 |
| CST-Household Influent-F7-R7 : CST-Household Influent-DF-DR | 0.973 |
| CST-Household Influent-F7-R7 : CST-Household Influent-F3-R3 | 0.913 |
| CST-Household Effluent-F3-R3 : CST-Household Effluent-DF-DR | 0.974 |
| CST-Household Effluent-F7-R7 : CST-Household Effluent-DF-DR | 0.93 |
| CST-Household Effluent-F7-R7 : CST-Household Effluent-F3-R3 | 0.83 |
| CST-Household Sludge-F3-R3 : CST-Household Sludge-DF-DR | 0.905 |
| CST-Household Sludge-F7-R7 : CST-Household Sludge-DF-DR | 0.995 |
| CST-Household Sludge-F7-R7 : CST-Household Sludge-F3-R3 | 0.859 |
| CST-Healthcare Effluent-F3-R3 : CST-Healthcare Effluent-DF-DR | NA |
| CST-Healthcare Effluent-F7-R7 : CST-Healthcare Effluent-DF-DR | NA |
| CST-Healthcare Effluent-F7-R7 : CST-Healthcare Effluent-F3-R3 | NA |
| CST-Healthcare Sludge-F3-R3 : CST-Healthcare Sludge-DF-DR | 0.827 |
| CST-Healthcare Sludge-F7-R7 : CST-Healthcare Sludge-DF-DR | 0.724 |
| CST-Healthcare Sludge-F7-R7 : CST-Healthcare Sludge-F3-R3 | 0.978 |
| SST-Household Effluent-F3-R3 : SST-Household Effluent-DF-DR | 0.843 |
| SST-Household Effluent-F7-R7 : SST-Household Effluent-DF-DR | 0.986 |
| SST-Household Effluent-F7-R7 : SST-Household Effluent-F3-R3 | 0.758 |
| SST-Household Sludge-F3-R3 : SST-Household Sludge-DF-DR | 0.908 |
| SST-Household Sludge-F7-R7 : SST-Household Sludge-DF-DR | 0.969 |
| SST-Household Sludge-F7-R7 : SST-Household Sludge-F3-R3 | 0.791 |

**Table S.****6:** Two-way Anova test between primer sets for the same sample types

CST, conventional septic tank; SST, solar septic tank; F, forward primer; R, reverse primer

**Table S.****7:** Summary statistics of the ASVs abundances per sample by MiSeq amplicon sequencing

| **Primer set** | **No of ASVs** | **ASV abundance summary statistics** | | | | | |
| --- | --- | --- | --- | --- | --- | --- | --- |
|  |  | **Minimum** | **1^st^ Quantile** | **Median** | **Mean** | **3^rd^ Quantile** | **Maximum** |
| DF-DR | 3 | 37801 | 47492 | 48890 | 51945 | 54056 | 113204 |
| F3-R3 | 4 | 40103 | 43406 | 45630 | 46602 | 48959 | 57196 |
| F7-R7 | 11 | 27723 | 34570 | 36653 | 36684 | 39880 | 45592 |

F, forward primer; R, reverse primer
